# Ankyrin-B is lipid-modified by S-palmitoylation to promote dendritic membrane scaffolding of voltage-gated sodium channel Nav1.2 in neurons

**DOI:** 10.1101/2022.06.01.494444

**Authors:** Julie P. Gupta, Paul M. Jenkins

## Abstract

Neuronal ankyrin-B is an intracellular scaffolding protein that plays multiple roles in the axon. By contrast, relatively little is known about the function of ankyrin-B in dendrites, where ankyrin-B is also localized in mature neurons. Recently, we showed that ankyrin-B acts as a scaffold for the voltage-gated sodium channel, Na_V_1.2, in dendrites of neocortical pyramidal neurons. How ankyrin-B is itself targeted to the dendritic membrane is not well understood. Here, we report that ankyrin-B is lipid-modified by S-palmitoylation to promote dendritic localization of Na_V_1.2. We identify the palmitoyl acyl transferase zDHHC17 as a key mediator of ankyrin-B palmitoylation in heterologous cells and in neurons. Additionally, we find that zDHHC17 regulates ankyrin-B protein levels independently of its S-acylation function through a conserved binding mechanism between the ANK repeat domain of zDHHC17 and the zDHHC ankyrin-repeat binding motif of ankyrin-B. We subsequently identify five cysteines in the N-terminal ankyrin repeat domain of ankyrin-B that are necessary for ankyrin-B palmitoylation. Mutation of these five cysteines to alanines not only abolishes ankyrin-B palmitoylation, but also prevents ankyrin-B from scaffolding Nav1.2 at dendritic membranes of neurons due to ankyrin-B’s inability to localize properly at dendrites. Thus, we show palmitoylation is critical for localization and function of ankyrin-B at dendrites. Strikingly, loss of ankyrin-B palmitoylation does not affect ankyrin-B-mediated axonal cargo transport of synaptic vesicle synaptotagmin-1 in neurons. This is the first demonstration of S-palmitoylation of ankyrin-B as an underlying mechanism required for ankyrin-B localization and function in scaffolding Nav1.2 at dendrites.

## INTRODUCTION

The ankyrin family of scaffolding proteins target and anchor membrane, cytoskeletal, and cytoplasmic proteins at specialized membrane domains to establish and maintain cell polarity and function in many vertebrate tissues (Bennett and Lorenzo, 2013). One member of the ankyrin family, ankyrin-B, encoded by the *ANK2* gene, localizes ion channels, transporters, structural proteins, as well as signaling molecules to specialized membranes in the heart, skeletal muscle, adipocytes, and brain (Ayalon et al., 2008;Ayalon et al., 2011;Lorenzo et al., 2014;Lorenzo and Bennett, 2017;Lorenzo, 2020;Sucharski et al., 2020)). In neurons, multiple axonal roles have been shown for ankyrin-B, including axonal cargo transport, scaffolding of cell adhesion molecules to repress axonal branching, and proper positioning of ankyrin-G at the axon initial segment (AIS) (Galiano et al., 2012;Lorenzo et al., 2014;Yang et al., 2019). By contrast, the role of ankyrin-B at dendritic membranes had been relatively understudied until recently, when ankyrin-B was shown to serve as an essential scaffold for voltage-gated sodium channels, Nav1.2, at dendrites to promote dendritic excitability and synaptic plasticity in neocortical pyramidal neurons (Nelson et al., 2022). Heterozygous loss of *Ank2* causes dramatic impairments in dendritic excitability and synaptic plasticity, and phenocopies effects of heterozygous loss of *Scn2a*, which encodes Nav1.2, suggesting ankyrin-B is critical for Nav1.2-mediated dendritic function (Spratt et al., 2019;Nelson et al., 2022).

*ANK2* is amongst the top genes linked to autism spectrum disorder (ASD) (Satterstrom et al., 2020). R2608fs, an ASD-associated variant in exon 37 of *ANK2*, which encodes the 440-kDa isoform of ankyrin-B, induces a frameshift that disrupts axonal scaffolding of the cell adhesion molecule L1CAM, causing alterations in axonal connectivity and behavior reminiscent of the deficits in communicative and social behaviors observed in human ASD patients (Yang et al., 2019). *SCN2A*, which encodes Nav1.2, is also a high-risk gene for ASD (Sanders et al., 2015;Ben-Shalom et al., 2017;Sanders et al., 2018). Nav1.2 and ankyrin-B interacting to promote dendritic excitability provides strong rationale that deficits in dendritic excitability may be a common point of convergence in ASD etiology. Although it is clear that Nav1.2 relies on ankyrin-B to properly target to dendritic membranes in neurons, how ankyrin-B itself reaches the dendritic membrane is unknown. Understanding how ankyrin-B targets to the dendritic membrane to mediate the localization and function of Nav1.2 may shed light onto ASD pathophysiology.

Another member of the ankyrin family, ankyrin-G, relies on the posttranslational modification, S-palmitoylation, to localize specifically to the lateral membrane of epithelial cells and to the axon initial segment (AIS) of neurons (He et al., 2012). S-palmitoylation is the covalent addition of a 16-carbon fatty acid to cysteine residues of proteins through the formation of a labile thioester bond, and is catalyzed by a family of 23 enzymes known as palmitoyl acyl transferases (zDHHC PATs) (Fukata and Fukata, 2010). Ankyrin-G is S-palmitoylated at a single cysteine in its N-terminal ankyrin repeat domain, Cys70 (He et al., 2012). Mutation of Cys70 to alanine (C70A) renders ankyrin-G palmitoylation-dead and subsequently incapable of associating with the lateral membrane of epithelial cells. Under normal circumstances, ankyrin-G assembles and maintains the localization of necessary components of the epithelial lateral membrane, such as βII-spectrin, to ensure membrane biogenesis and cell polarity. However, palmitoylation-dead C70A ankyrin-G is incapable of clustering itself as well as βII-spectrin to the epithelial lateral membrane, resulting in loss of membrane biogenesis and loss of membrane height (He et al., 2012;He et al., 2014). In cultured hippocampal neurons, rescue of Cre-dependent deletion of ankyrin-G at the AIS with C70A ankyrin-G prevents ankyrin-G from specifically clustering at the AIS and instead distributes ankyrin-G in a non-polarized manner throughout neuronal compartments (He et al., 2012;Tseng et al., 2015). Ankyrin-G is palmitoylated by two zDHHC PATs, zDHHC5 and zDHHC8 (He et al., 2014). Knockdown of zDHHC5 and zDHHC8 abolishes ankyrin-G palmitoylation and phenocopies the loss of membrane height and biogenesis observed with C70A ankyrin-G (He et al., 2014). Taken together, these data demonstrate that ankyrin-G S-palmitoylation is required for ankyrin-G localization and function at the lateral membrane of epithelial cells as well as at the neuronal AIS. Whether ankyrin-B is also palmitoylated as a mechanism to regulate its plasma membrane localization and scaffolding of Nav1.2 at dendrites remains unclear.

Here, we asked if ankyrin-B was S-palmitoylated and whether palmitoylation could regulate ankyrin-B localization and function in neurons. We find that ankyrin-B is indeed palmitoylated in whole mouse brain and in neurons, identifying zDHHC17 as a key mediator of ankyrin-B palmitoylation in heterologous cells and in neurons. We also demonstrate that zDHHC17 regulates ankyrin-B protein levels in a palmitoylation-independent manner through a conserved binding mechanism between the ANK repeat of zDHHC17 and the zDHHC ankyrin-repeat binding motif (zDABM) of ankyrin-B. We identify five cysteines as possible ankyrin-B palmitoylation sites. Mutation of these five cysteines in the N-terminal ANK repeat domain of ankyrin-B renders ankyrin-B palmitoylation-dead. While ankyrin-B palmitoylation does not regulate its ability to mediate axonal cargo function, ankyrin-B palmitoylation is required to target Nav1.2 to dendrites in neocortical pyramidal neurons. These data provide evidence for two pools of ankyrin-B in neurons, one palmitoylation-dependent pool at dendrites, and one palmitoylation-independent pool in vesicles at the axon. This work highlights novel mechanisms regulating ankyrin-B localization and function in neurons that may provide insights into *ANK2-* associated ASD.

## RESULTS

### Ankyrin-B is S-palmitoylated in whole mouse brain and in neurons

The homologous family member of ankyrin-B, ankyrin-G, is S-palmitoylated to drive ankyrin-G localization and function in epithelial cells and in neurons (He et al., 2012). Ankyrin-G and ankyrin-B share significant homology across their shared domains, with 74% homology shared in their N-terminal ankyrin repeat domain, where ankyrin-G’s palmitoylated cysteine resides (Mohler et al., 2002). Given such high homology, we asked whether ankyrin-B is also S-palmitoylated and if so, whether S-palmitoylation regulates ankyrin-B’s diverse functions in neurons. We used the Acyl Resin Assisted Capture (Acyl-RAC) assay to assess steady-state palmitoylation of ankyrin-B (previously described (Bouza et al., 2020)). To validate this assay in our laboratory, we confirmed previous results demonstrating ankyrin-G is S-palmitoylated (He et al., 2012;Tseng et al., 2015). Using C57Bl/6J adult mouse whole-brain lysates subjected to Acyl-RAC, we observed that all three neuronal isoforms of ankyrin-G, 190-, 270-, and 480-kDa ankyrin-G, are S-palmitoylated, as demonstrated by hydroxylamine-dependent binding of endogenous ankyrin-G to sepharose beads (“+HA” lane) (Figure 1A, B). We also observed that the 220-kDa and 440-kDa isoforms of ankyrin-B are S-palmitoylated in whole mouse brain, as evidenced by the presence of hydroxylamine-dependent ankyrin-B signal (Figure 1A, C). Given our interest in understanding whether palmitoylation regulates the diverse functions of ankyrin-B in neurons, we asked whether ankyrin-B is also palmitoylated in cultured hippocampal neurons from C57Bl/6J mice. Cultured hippocampal neurons represent a rational model from which to test ankyrin-B palmitoylation given that the axonal cargo transport function of ankyrin-B was previously established in hippocampal neurons (Lorenzo et al., 2014). Indeed, we observed palmitoylation of both 220- and 440-kDa isoforms of ankyrin-B in cultured hippocampal neurons by Acyl-RAC, as shown by the hydroxylamine-dependent ankyrin-B signal (Figure 1D). Taken together, these data provide the first evidence that ankyrin-B is S-palmitoylated in whole mouse brain and in cultured neurons, implicating palmitoylation as a potential mechanism underlying ankyrin-B localization and function in neurons.

**Figure 1.**
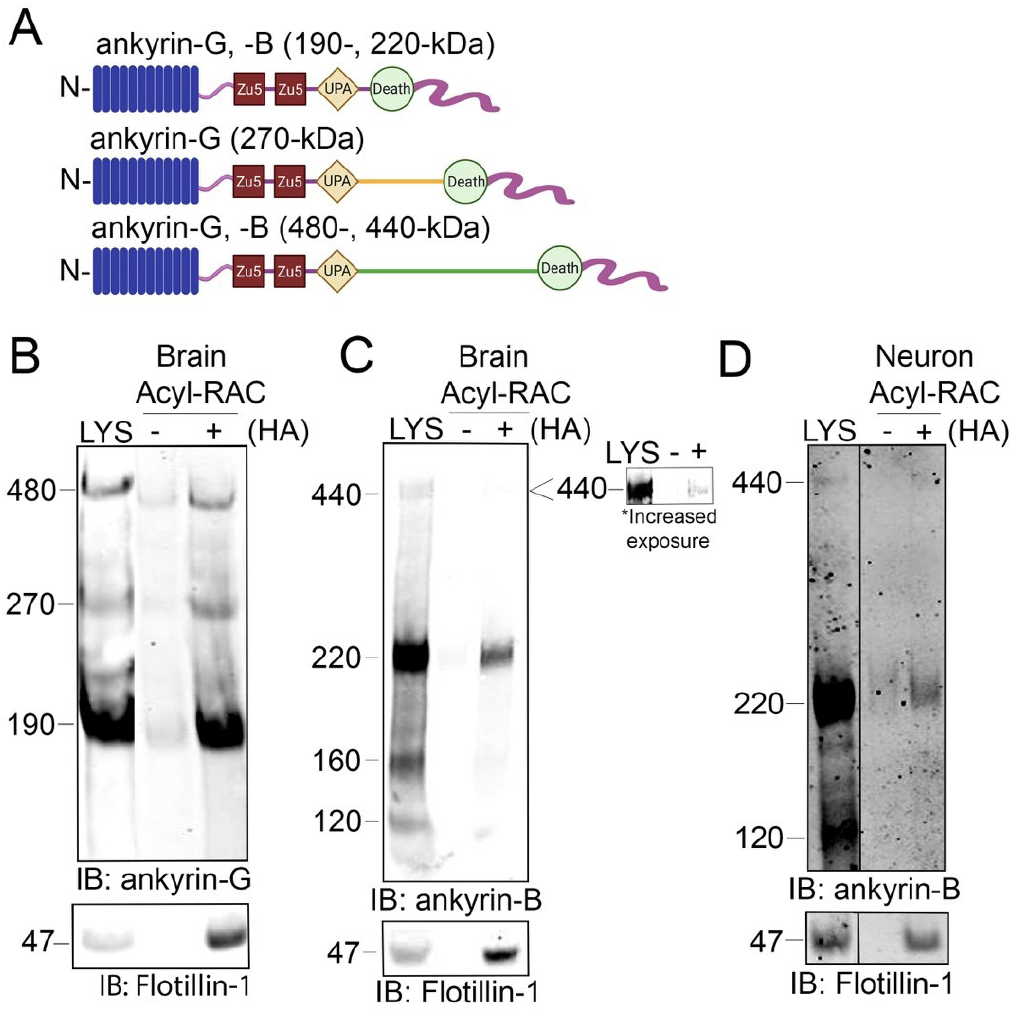
Ankyrin-B is S-palmitoylated in whole mouse brain and in cultured neurons. **A:** Schematic of neuronal ankyrin-G and ankyrin-B, which share highly homologous domains: the N-terminal ankyrin repeat domain (blue), two ZU5 spectrin binding domains (red), a UPA domain (yellow), a death domain (light green), and a highly divergent unstructured C-terminal regulatory domain (pink). The giant isoforms (480-kDa ankyrin-G and 440-kDa ankyrin-B) have a large single exon (orange and dark green) inserted between their respective UPA and death domains. **B:** Whole brains from C57Bl/6J mice were processed for the Acyl-RAC assay to detect S-palmitoylation. S-palmitoylation of all neuronal isoforms of ankyrin-G (190-, 270-, and 480-kDa) is detected in whole brain from C57Bl/6J mice using an antibody against endogenous ankyrin-G, as demonstrated by the anti-ankyrin-G signal in the ‘+HA’ lane, compared to the minimal background signal in the negative control ‘-HA’ lane. Flotillin-1 is used as a positive control for the Acyl-RAC assay (N=3). **C:** S-palmitoylation of the known neuronal isoforms of ankyrin-B (220- and 440-kDa) is detected by Acyl-RAC in whole brain from C57Bl/6J mice using an antibody against endogenous ankyrin-B (N=3). **D:** DIV17 cultured hippocampal neurons from C57Bl/6J mice were subjected to the Acyl-RAC assay. S-palmitoylation of the 220-kDa and 440-kDa isoforms of ankyrin-B is detected in cultured hippocampal neurons using an antibody against endogenous ankyrin-B (N=3). All conditions shown are run on the same blot; black line delineates spliced portion of the gel containing a condition not relevant to this figure. IB, immunoblotting

### Ankyrin-B expression and S-palmitoylation are increased by the palmitoyl acyltransferase zDHHC17 in heterologous cells

Multiple studies have shown that the palmitoyl acyltransferase (PAT) zDHHC17 binds tightly to its substrates in order to palmitoylate them (Lemonidis et al., 2014;Verardi et al., 2017). The N-terminal ANK repeat (AR) domain of zDHHC17 recognizes a conserved zDHHC AR-binding motif (zDABM), (VIAP)(VIT)XXQP (where X is any amino acid) in its substrates (Lemonidis et al., 2014;Lemonidis et al., 2015;Verardi et al., 2017). Interestingly, a previous study identified the zDABM domain in the ankyrin family, including in ankyrin-B, using a peptide array screening approach (Figure 2A) (Lemonidis et al., 2017). This finding provided rationale for investigating the role of zDHHC17 in ankyrin-B palmitoylation. We subjected lysates from HEK293T cells transiently transfected with HA-tagged zDHHC17 and ankyrin-B-GFP to Acyl-RAC and found that zDHHC17 significantly enhanced both the palmitoylation and protein levels of ankyrin-B-GFP, compared to ankyrin-B-GFP in the absence of any exogenous PATs (Figure 2B-E). The presence of GFP signal in the negative control “-HA” lane is likely due to incomplete blocking of all free cysteines in ankyrin-B-GFP, presumably due to the GFP tag, which has numerous cysteines embedded in stable secondary structure that our lysis conditions may not disrupt completely. However, our quantification accounts for this background signal by subtracting any signal in the “-HA” lane from the “+HA” signal. To ensure that the increase in palmitoylation and expression levels of ankyrin-B-GFP seen with zDHHC17 were PAT-specific and not a function of overexpression, we screened the zDHHC library consisting of all 23 PATs (Figure S1A-D). Using Acyl-RAC, we observed most zDHHC PATs did not enhance the palmitoylation nor the expression levels of ankyrin-B-GFP, compared to ankyrin-B-GFP alone, suggesting that the assay is able to detect PAT-dependent differences in palmitoylation and protein expression. Consistent with our results from Figure 3.2, our PAT screen revealed zDHHC17 was the only exogenous PAT able to significantly enhance the palmitoylation of ankyrin-B, compared to ankyrin-B alone (Figure S1A, B). zDHHC17 was also the only PAT that significantly increased ankyrin-B protein expression, compared to ankyrin-B alone (Figure S1C, D). To test whether the increase in ankyrin-B-GFP protein levels seen in the presence of zDHHC17 was dependent on palmitoylation, we transiently transfected ankyrin-B-GFP with a catalytically-dead version of HA-tagged zDHHC17 (zDHHA17) in HEK293T cells and subjected the lysates to Acyl-RAC. zDHHA17 retains its recognition site for substrates (ANK repeat domain) but loses its enzymatic activity as the cysteine of the catalytic DHHC motif (C467) is mutated to alanine (Locatelli et al., 2020). We observed that palmitoylation of ankyrin-B-GFP was completely abolished in the presence of enzymatically-dead zDHHA17, as compared to that with WT zDHHC17 (Figure 2B-D). In fact, there was no significant difference in ankyrin-B-GFP palmitoylation between ankyrin-B-GFP alone and ankyrin-B-GFP co-expressed with zDHHA17 (Figure 2B-D). Thus, the increase in ankyrin-B-GFP palmitoylation observed upon co-expression of WT zDHHC17 is indeed driven by the catalytic activity of zDHHC17. Notably, however, ankyrin-B-GFP levels were still increased by zDHHA17, similar to what we observed with WT zDHHC17 (Figure 2B, E). These results suggest that the observed increase in ankyrin-B-GFP protein levels in the presence of zDHHC17 is not palmitoylation-dependent and may instead be due to recognition of ankyrin-B-GFP by zDHHC17. To test whether the enhancement in ankyrin-B protein levels are dependent on zDHHC17 recognition, we co-expressed a version of zDHHC17 whose substrate binding site is abolished, the HA-tagged N100A zDHHC17. N100A zDHHC17 is a well-characterized ANK repeat mutant that fails to recognize zDABM-containing substrates (Verardi et al., 2017;Niu et al., 2020). Notably, ankyrin-B-GFP protein levels were reduced approximately three fold in the presence of N100A zDHHC17 compared to that with WT or zDHHA17 (Figure 2B, E), suggesting that zDHHC17 recognition of ankyrin-B regulates ankyrin-B protein expression in heterologous cells. Surprisingly, ankyrin-B protein stability was unchanged in the presence of WT or N100A zDHHC17, compared to ankyrin-B alone by cycloheximide chase assay, suggesting that zDHHC17 is not regulating ankyrin-B expression by conferring stability in heterologous cells (Figure S2A, B). Future studies will need to investigate the mechanisms underlying zDHHC17-dependent increases in ankyrin-B protein expression in heterologous cells. Given that zDHHC17 often requires direct interaction with its substrates to palmitoylate them (Lemonidis et al., 2017), we hypothesized that the N100A mutation in zDHHC17 would also prevent palmitoylation. As predicted, loss of zDHHC17-ankyrin-B recognition induced by the N100A mutation in zDHHC17 or zDHHA17 results in drastically reduced palmitoylation levels compared to that with WT zDHHC17 (Figure 2B-D). These results are consistent with the canonical zDHHC17 AR-substrate zDABM binding paradigm necessary for palmitoylation, and confirms the findings of the peptide array screen that showed ankyrin-B has a zDABM domain (Lemonidis et al., 2017).

**Figure 2.**
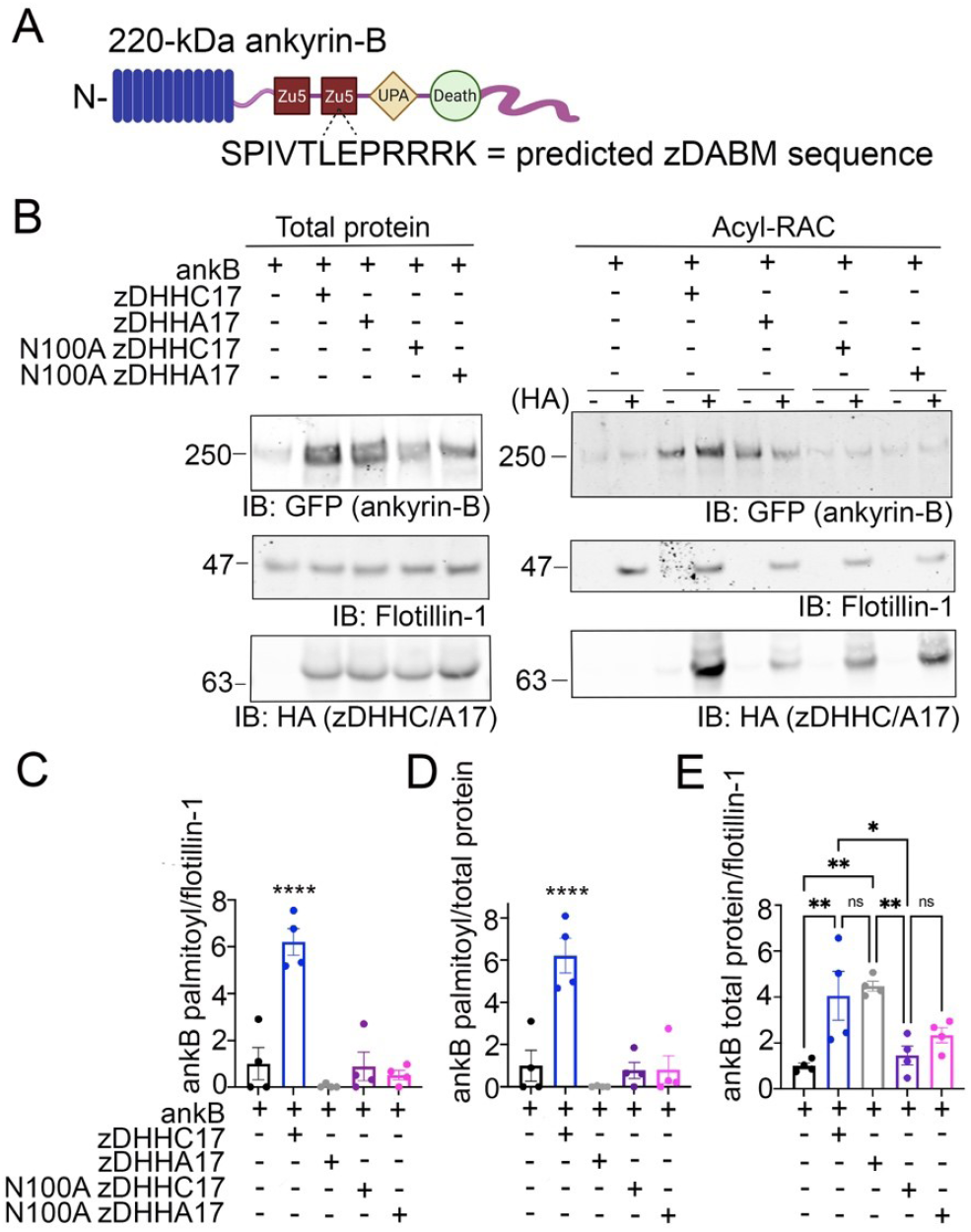
zDHHC17 recognizes the zDABM domain of ankyrin-B to regulate ankyrin-B protein expression and palmitoylation in HEK293T cells. **A:** Schematic of ankyrin-B with the predicted zDHHC ANK repeat binding motif sequence (zDHHC17 recognition site, predicted from [27]) located in the second ZU5 domain. **B:** Representative western blot showing total protein levels of ankyrin-B-GFP (left) and palmitoylation levels of ankyrin-B-GFP (right) from lysates of HEK293T cells transiently transfected with WT ankyrin-B-GFP alone or together with WT zDHHC17, enzymatically-dead zDHHA17, N100A zDHHC17, or N100A zDHHA17 subjected to the Acyl-RAC assay. Ankyrin-B-GFP protein and palmitoylation levels are detected with an antibody against GFP. Palmitoylation of ankyrin-B-GFP is strongly detected in a hydroxylamine-dependent manner (‘+HA’ lane), compared to the background signal in the negative control ‘-HA’ lane. Co-expression of zDHHC17 increases both the protein levels (left) and palmitoylation levels (right) of ankyrin-B-GFP, compared to ankyrin-B expressed alone. WT ankyrin-B-GFP with zDHHA17, N100A zDHH17, or N100A zDHHA17 is not palmitoylated, compared to when ankyrin-B-GFP is co-expressed with zDHHC17. Ankyrin-B-GFP protein levels are unaffected by co-expression of zDHHA17, but are drastically reduced with co-expression of N100A zDHHC17 or N100A zDHHA17. **C:** Quantified S-palmitoylation levels of ankyrin-B-GFP from N=4 independent replicates per condition from A. ****P<0.0001 relative to ankyrin-B alone; one way ANOVA with Tukey’s post-hoc test. For each condition, the palmitoylation signal was calculated by subtracting the -HA lane signal from that of the +HA lane and normalized to the flotillin-1 signal from the +HA lane, and normalized again to the average of ankyrin-B alone signal to get the relative fold change in ankyrin-B-GFP palmitoylation. **D:** Quantified ankyrin-B-GFP S-palmitoylation levels normalized to total ankyrin-B protein levels for each condition, relative to ankyrin-B alone palmitoylation levels. Data from N=4 independent replicates per condition from A. ****P<0.0001 relative to ankyrin-B alone; one way ANOVA with Tukey’s post-hoc test. **E:** Quantified ankyrin-B-GFP protein levels normalized to total flotillin-1 levels, relative to ankyrin-B alone protein levels. Data from N=4 independent replicates per condition from A. **P<0.01, *P<0.05; one way ANOVA with Tukey’s post-hoc test.

### zDHHC17 is required to palmitoylate endogenous ankyrin-B in cultured hippocampal neurons

Our findings using overexpression in heterologous cells suggest that zDHHC17 is a strong candidate for ankyrin-B palmitoylation. To address whether zDHHC17 palmitoylates endogenous ankyrin-B, we used cultured hippocampal neurons from floxed *Zdhhc17* mice (Sanders et al., 2016) that were transduced with either a β-galactose control adenovirus or a Cre-recombinase adenovirus before lysates were either subjected to RT-PCR analysis to probe for extent of *Zdhhc17* deletion or the Acyl-RAC assay to probe for effects on ankyrin-B palmitoylation. RT-PCR analysis revealed a 95% reduction of *Zdhhc17* mRNA levels ten days post-transduction of Cre adenovirus in floxed *Zdhhc17* hippocampal neurons, compared to neurons treated with β-galactose control adenovirus (Figure 3A). Acyl-RAC revealed an 80% loss of ankyrin-B palmitoylation in Cre-treated floxed *Zdhhc17* hippocampal neurons, compared to that with the control virus (Figure 3B-D). Results from Figure 2 demonstrating that zDHHC17 recognizing ankyrin-B is sufficient to increase protein levels led us to hypothesize that loss of endogenous zDHHC17 would reduce endogenous ankyrin-B protein levels. However, there was no significant difference in ankyrin-B protein levels between the Cre virus-treated and the control virus-treated neurons (Figure 3E), suggesting that neurons may utilize mechanisms independent of zDHHC17 to regulate ankyrin-B protein levels, unlike in heterologous cells. Taken together, these data demonstrate zDHHC17 is a major regulator of ankyrin-B palmitoylation in neurons.

**Figure 3.**
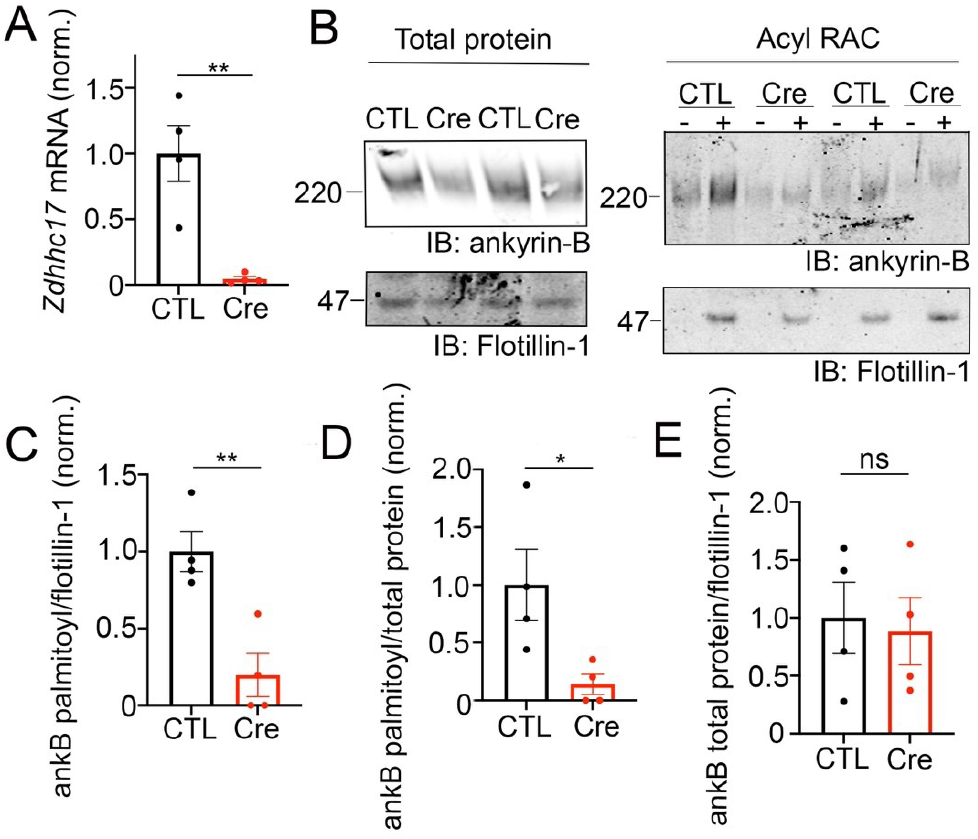
zDHHC17 recognizes the zDABM domain of ankyrin-B to regulate ankyrin-B protein expression and palmitoylation in HEK293T cells. **A:** Zdhhc17 mRNA levels in cultured hippocampal from Zdhhc17 mice transduced with either a β-galactose control adenovirus or Cre-recombinase adenovirus for 10 days prior to sample collection for RT-PCR analysis (n=4). Zdhhc17 mRNA was decreased by ~95% in the Cre-recombinase condition compared to control. **P<0.01; Student’s t-test (unpaired). **B:** Representative western blot showing total protein levels of endogenous ankyrin-B (left) and palmitoylation levels of endogenous ankyrin-B (right) from lysates of floxed Zdhhc17 hippocampal neurons transduced with either a β-galactose control adenovirus or Cre-recombinase adenovirus for 10 days before Acyl-RAC processing (N=4). Both protein (left) and palmitoylation (right) levels of endogenous ankyrin-B are immunoblotted with an antibody against ankyrin-B for all conditions. Palmitoylation of ankyrin-B is strongly detected in a hydroxylamine dependent manner (‘+HA’ lane) from cultures infected with the control virus, but is greatly reduced in cultures infected with the Cre virus, as demonstrated by the lower ankyrin-B signal intensity in the ‘+HA’ lane. **C:** Quantified endogenous ankyrin-B S-palmitoylation normalized to flotillin-1 levels from the Acyl-RAC fractions and further normalized to control virus. Data from N=4 independent replicates per condition from A. **P<0.01; Student’s t-test (unpaired). **D:** Quantified endogenous ankyrin-B S-palmitoylation normalized to total ankyrin-B protein levels for each condition, and further normalized to control virus. Data from N=4 independent replicates per condition from A. **P<0.05; Student’s t-test (unpaired). **E:**Quantified endogenous ankyrin-B levels normalized to total flotillin levels for each condition, and further normalized to control virus. Data from N=4 independent replicates per condition from A. not significant (ns); Student’s t-test (unpaired).

### Ankyrin-B is S-palmitoylated at multiple cysteines in its N-terminal ankyrin repeat domain

The palmitoylation sites within ankyrin-B are unknown. Ankyrin-B’s homologous family member, ankyrin-G, is S-palmitoylated at a single cysteine in its N-terminal ankyrin repeat domain, Cys70 (He et al., 2012). Cys70 is conserved among all three human ankyrin members and vertebrate ankyrins (He et al., 2012), with ankyrin-B possessing a homologous cysteine at Cys60 (Figure 4A). Given the conservation of this amino acid, we hypothesized that Cys60 in ankyrin-B is the principal palmitoylated cysteine. Using site-directed mutagenesis, we engineered a cDNA construct in which Cys60 in ankyrin-B was converted to an alanine (C60A ankyrin-B-GFP). The C60A ankyrin-B-GFP mutant construct was transiently transfected with HA-tagged zDHHC17 into HEK293T cells, and lysates were subjected to Acyl-RAC to test the mutant’s effects on ankyrin-B palmitoylation. The C60A ankyrin-B mutant lead to a 60% reduction in ankyrin-B palmitoylation, compared to WT ankyrin-B, despite efficient expression of C60A ankyrin-B-GFP (Figure 4B, C). These data suggest that there are additional palmitoylated cysteine sites in ankyrin-B beyond Cys60.

**Figure 4.**
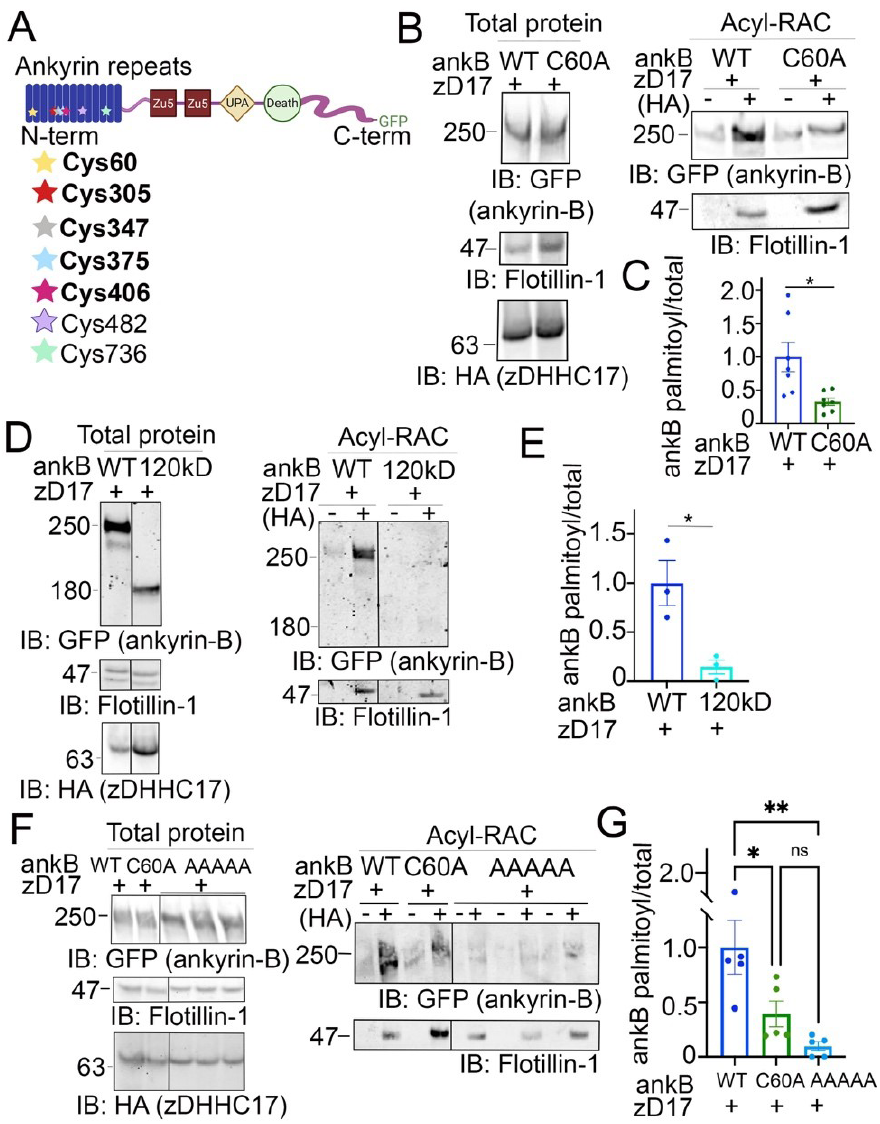
Identification of ankyrin-B S-palmitoylated cysteine sites. **A:** Schematic of the domains in ankyrin-B and the cysteine residues harbored in the N-terminal ankyrin repeat domain, which are candidates palmitoylation sites. Bolded cysteine residues are the cysteine residues that were substituted to alanine to generate the palmitoylation-dead ankyrin-B mutant. **B.:**Representative western blot showing total protein levels of ankyrin-B-GFP (left) and palmitoylation levels of ankyrin-B-GFP (right) from lysates of HEK293T cells transiently co-transfected with WT ankyrin-B-GFP or C60A ankyrin-B-GFP and zDHHC17 processed for the Acyl-RAC assay (N=7). S-palmitoylation of C60A ankyrin-B-GFP is greatly reduced, as evidenced by the lower GFP signal in the ‘+HA’ lane compared to that of the WT ankyrin-B-GFP. **C:** Quantified ankyrin-B-GFP S-palmitoylation levels normalized to total ankyrin-B protein levels for each condition, relative to palmitoylation levels of WT ankyrin-B-GFP co-expressed with zDHHC17. Data from N=7 independent replicates per condition from A. *P=0.0119; Student’s t-test (unpaired). **D:** Representative western blot showing total protein levels of ankyrin-B-GFP (left) and palmitoylation levels of ankyrin-B-GFP (right) from lysates of HEK293T cells transiently co-transfected with WT ankyrin-B-GFP or 120-kDa ankyrin-B-GFP and zDHHC17 processed for the Acyl-RAC assay (N=3). All conditions shown are run on the same blot; black line delineates spliced portion of the gel containing a condition not relevant to this figure. S-palmitoylation of 120-kDa ankyrin-B-GFP is not detected using an antibody against GFP, as shown by the absence of anti-GFP signal in the ‘+HA’ lane, compared to that when WT ankyrin-B-GFP is co-expressed with zDHHC17. **E:** Quantified ankyrin-B-GFP S-palmitoylation levels normalized to total ankyrin-B protein levels for each condition, relative to palmitoylation levels of WT ankyrin-B-GFP co-expressed with zDHHC17. Data from N=3 independent replicates per condition from D. *P=0.0241; Student’s t-test (unpaired). **F:** Representative western blot showing total protein levels of ankyrin-B-GFP (left) and palmitoylation levels of ankyrin-B-GFP (right) from lysates of HEK293T cells transiently co-transfected with WT ankyrin-B-GFP, C60A ankyrin-B-GFP, or AAAAA-ankyrin-B-GFP and zDHHC17 processed for the Acyl-RAC assay (N=5). All conditions shown are run on the same blot; black line delineates spliced portion of the gel containing a condition not relevant to this figure. Compared to WT ankyrin-B-GFP and C60A ankyrin-B-GFP, S-palmitoylation of AAAAA-ankyrin-B-GFP is almost completely abolished. **G:** Quantified ankyrin-B-GFP S-palmitoylation levels normalized to total ankyrin-B protein levels for each condition, relative to palmitoylation levels of WT ankyrin-B-GFP co-expressed with zDHHC17. Data from N=5 independent replicates per condition from F. **P<0.01, *P<0.05; one way ANOVA with Tukey’s post-hoc test.

Ankyrin-B has 26 cysteines, seven of which are harbored in the N-terminal ankyrin repeat domain, a domain necessary for ankyrins to interact with their membrane-associated partners (Figure 4A) (Bennett and Lorenzo, 2013). To narrow down on additional palmitoylation sites in ankyrin-B, we asked whether palmitoylation sites were restricted to the N-terminal ankyrin repeat domain of ankyrin-B. Acyl-RAC on lysates from HEK293T cells transiently transfected with a truncated version of ankyrin-B which completely lacks the ankyrin repeat domain (120-kDa ankyrin-B-GFP) as well as zDHHC17 demonstrated an almost complete reduction of ankyrin-B-GFP palmitoylation, compared to WT ankyrin-B-GFP, despite efficient expression of 120-kDa ankyrin-B-GFP (Figure 4D, E). These data suggest that additional palmitoylation sites of ankyrin-B are located within the ankyrin repeat domain.

To determine which cysteines may be additional palmitoylation sites within the ankyrin-B ankyrin repeat domain, we employed mass spectrometry to identify peptides with S-palmitoylated cysteine sites in ankyrin-B. Lysates from HEK293T cells transfected with ankyrin-B-GFP and zDHHC17-HA were processed for the Acyl-RAC assay, digested, and analyzed by nano Liquid Chromatography tandem Mass Spectrometry (LC-MS/MS). With this approach, we identified five ankyrin-B peptides containing Cys60, Cys305, Cys347, Cys375, and Cys406 (Figure 4A (bolded), Figure S3A). We did not detect peptides containing Cys482 and Cys736, likely indicating that these cysteines were not palmitoylated (Figure S3A). To confirm that Cys482 and Cys736 were not palmitoylation sites in ankyrin-B, we engineered cDNA constructs containing a cysteine-to-alanine mutation at C482A and another at C736A in ankyrin-B-GFP. C482A ankyrin-B-GFP and C736A ankyrin-B-GFP were individually expressed with zDHHC17-HA in HEK293T cells and processed for the Acyl-RAC assay to test their individual effects on ankyrin-B palmitoylation. Indeed, co-expression of zDHHC17-HA and either C482A ankyrin-B-GFP or C736A ankyrin-B-GFP did not affect ankyrin-B palmitoylation, compared to WT ankyrin-B (Figure S3B, C).

To further investigate the functional role of the remaining four candidate palmitoylation sites in ankyrin-B identified in the mass spectrometry study (C305, C347, C375, C406) (Figure 4A), we constructed a mutant version of ankyrin-B with the five cysteine residues, Cys60 included, substituted to alanine residues using site-directed mutagenesis. This mutant GFP-tagged ankyrin-B construct is referred to as AAAAA-ankyrin-B-GFP (or A5B), and when co-expressed with zDHHC17, demonstrated virtually no ability to be palmitoylated compared to WT ankyrin-B-GFP, despite efficient expression of this mutant (Figure 4F, G). This is shown by the loss of hydroxylamine-dependent GFP signal in the presence of AAAAA-ankyrin-B-GFP, compared to WT and C60A ankyrin-B-GFP. These data suggest that ankyrin-B is palmitoylated at multiple cysteine sites harbored in its N-terminal ankyrin-repeat domain. Furthermore, the palmitoylation-dead ankyrin-B mutant, AAAAA-ankyrin-B-GFP, represents a novel tool for investigating the functional consequences of ankyrin-B palmitoylation on downstream ankyrin-B localization and function.

### S-palmitoylation does not regulate ankyrin-B-mediated axonal cargo transport in hippocampal neurons

Deletion of *Ank2*, which encodes ankyrin-B, leads to impaired organelle transport in axons of cultured hippocampal neurons (Lorenzo et al., 2014). Here, we asked whether palmitoylation could regulate the axonal transport function of ankyrin-B. To study the effects of the palmitoylation-dead form of ankyrin-B on an ankyrin-B-null background, we used cultured hippocampal neurons from floxed *Ank2* mice to knock out ankyrin-B with transfection of Cre recombinase and simultaneously rescued with either wild-type (WT) 220-kDa ankyrin-B-GFP or palmitoylation-dead AAAAA-ankyrin-B-GFP (A5B) cDNA. Transfection of Cre-recombinase and rescue cDNA at 2 days in vitro (DIV) and imaging by time-lapse video microscopy 48 hours later allowed for observation of optimal motility of synaptic vesicle protein tdTomato-tagged synaptotagmin-1 (Syt1-tdTomato) along the axon in real time. Syt1-TdTomato moved bidirectionally along the axon with high processivity in control neurons only transfected with Syt1-tdTomato (Figure 5A, B), as observed previously (Lorenzo et al., 2014). In neurons transfected with Syt1-tdTomato and Cre recombinase-2A-BFP to knock out ankyrin-B, we observed an increased percentage of stationary or trapped synaptic vesicles in axonal swellings along the axons (Figure 5B). Additionally, the Syt1-tdTomato vesicles that still retained motility in the Cre-knockout condition exhibited slower velocities and traveled shorter distances bidirectionally along the axons, compared to controls (Figure 5A, B), which recapitulated the organelle transport deficits observed in the AnkB-/- mice (Lorenzo et al., 2014). The retrograde velocity deficits of Syt1-tdTomato particles observed upon deletion of ankyrin-B (Cre-2A-BFP) were partially rescued with addition of WT 220-kDa ankyrin-B-GFP or palmitoylation-dead AAAAA-ankyrin-B-GFP, whereas neither WT or AAAAA-ankyrin-B-GFP rescued the anterograde velocity deficits of Syt1 particles induced by deletion of ankyrin-B (Figure 5A, B). The retrograde run length deficits of Syt1-tdTomato particles in the absence of ankyrin-B (Cre-2A-BFP) were fully rescued with addition of WT ankyrin-B-GFP or palmitoylation-dead AAAAA-ankyrin-B-GFP, whereas neither WT or AAAAA-ankyrin-B-GFP rescued anterograde run length deficits of Syt1 particles induced by deletion of ankyrin-B (Figure 5A, B). Longer expression times may be necessary to observe rescue of anterograde velocity or run length deficits with WT ankyrin-B-GFP. Addition of WT or AAAAA-ankyrin-B-GFP also rescued the increase in Syt1-tdTomato stationary particles induced by deletion of ankyrin-B (Cre-2A-BFP) (Figure 5B). Furthermore, we did not observe any significant differences in the speed, distance, or relative mobility of WT or palmitoylation-dead AAAAA-ankyrin-B-GFP particles (Figure 5C, D). Taken together, these results suggest palmitoylation-dead ankyrin-B is capable of mediating axonal cargo transport of Syt1 similar to WT ankyrin-B. Thus, ankyrin-B-mediated cargo transport of synaptic vesicles like Syt1 is not dependent on ankyrin-B palmitoylation.

**Figure 5.**
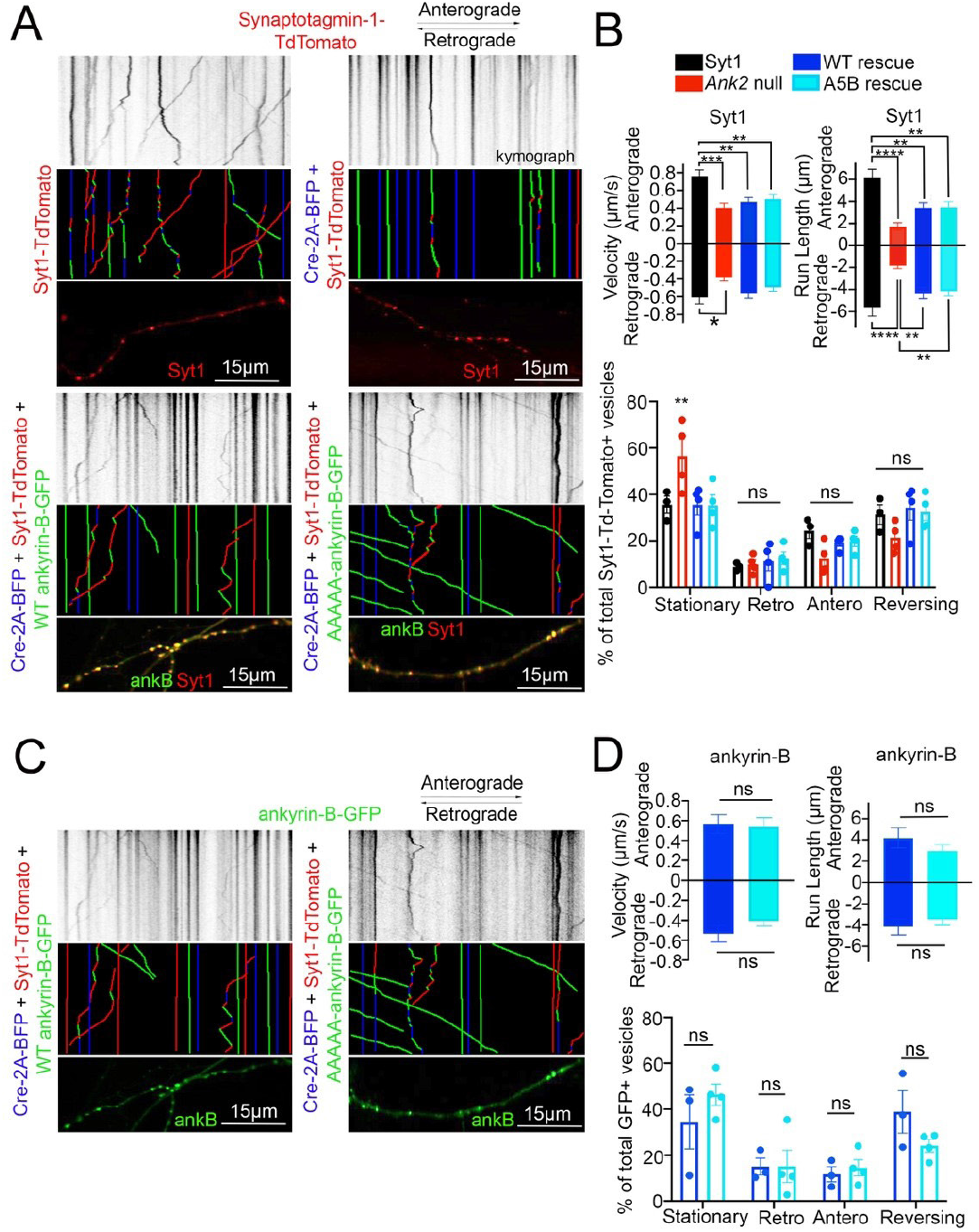
Loss of ankyrin-B palmitoylation does not affect axonal cargo transport of synaptic vesicle protein Synaptotagmin-1 (Syt1). **A:** (Top) Representative kymographs of Td-tomato-tagged Syt1 particle movement along DIV4 axons of floxed Ank2 hippocampal neurons in the absence or presence of Cre-2A-BFP to knock-out ankyrin-B and WT or palmitoylation-dead AAAAA-ankyrin-B-GFP as rescue plasmids. (Middle) Corresponding color-coded trajectories for Td-Tomato-positive particles demonstrating static vesicles (blue), anterograde-moving particles (green), and retrograde-moving particles (red). (Bottom) Puncta distribution of Syt1-TdTomato vesicles along axons (region of axon used to generate kymographs). **B:** Axonal velocity and run length for Syt1 particles. Data shown is from one representative experiment out of 4 independent repeats. N=4-8 axons per condition (n=88-154 particles per condition). Data represent means ± SEM. **P<0.01, ***P<0.001, ****P<0.0001. For velocity and run lengths analyses, one way ANOVA with Tukey’s post-hoc test were performed. For % motility analysis, two-way ANOVA with multiple comparisons and Tukey’s post-hoc was performed. **C:** (Top) Representative kymographs of GFP-tagged ankyrin-B particle movement along DIV4 axons of floxed Ank2 hippocampal neurons in the absence or presence of Cre-2A-BFP to knock-out ankyrin-B and WT or palmitoylation-dead AAAAA-ankyrin-B-GFP as rescue plasmids. (Middle) Corresponding color-coded trajectories for GFP-positive particles demonstrating static vesicles (blue), anterograde-moving particles (green), and retrograde-moving particles (red). (Bottom) Puncta distribution of WT or AAAAA-ankyrin-B-GFP vesicles along axons (region of axon used to generate kymographs). **D**: Axonal velocity and run length for ankyrin-B particles. Data shown is from one representative experiment out of 4 independent repeats. N=3-7 axons per condition (n=38-76 particles per condition). Data represent means ± SEM. non-significant (ns). For velocity and run lengths analyses, Student’s t-tests were performed. For % motility analysis, two-way ANOVA with multiple comparisons and Tukey’s post-hoc was performed.

### S-palmitoylation is required for ankyrin-B-mediated scaffolding of Nav1.2 at the dendritic membranes of neocortical pyramidal neurons

Given the lack of effect in axonal transport, we asked whether palmitoylation of ankyrin-B could serve other functions. Recently, we showed that ankyrin-B scaffolds the voltage-gated sodium channel Nav1.2 at the dendritic membrane to promote dendritic excitability and synaptic function (Nelson et al., 2022). Here, we asked whether palmitoylation was required for ankyrin-B to scaffold Nav1.2 at the dendritic membrane. Given that the dendritic deficits observed upon heterozygous loss of ankyrin-B in vivo were observed in the prefrontal layer 5b of mouse cortex and loss of Nav1.2 dendritic localization due to ankyrin-B deletion were observed in neocortical neuron cultures (Nelson et al., 2022), we investigated the role of ankyrin-B palmitoylation for Nav1.2 scaffolding in neocortical cultures. To validate that Nav1.2 localized properly at the AIS, where it is canonically scaffolded by ankyrin-G, as well as at dendritic membranes, we transfected DIV2 floxed *Ank2* neocortical neurons with Na_V_1.2-3x FLAG, fixed at DIV21 (when the NaV1.2 AIS-to-dendrite shift has occurred (Spratt et al., 2019)), and stained with antibodies against FLAG and ankyrin-B. We observed intense Nav1.2-3x FLAG staining at the AIS, consistent with Nav1.2’s role at the AIS in mature neurons (referenced by the white star in Figure 6A), and Nav1.2-3x FLAG staining at dendritic membranes, as evidenced by the “railroad track” footprint that overlaps with endogenous ankyrin-B staining (Figure 6A). To validate that deletion of ankyrin-B in neocortical neurons results in loss of Nav1.2-3x FLAG dendritic localization (Nelson et al., 2022), we transfected DIV2 floxed *Ank2* neocortical neurons with NaV1.2-3x FLAG and Cre-2A-BFP to knockout ankyrin-B. As expected, we observed nearly complete reduction in Nav1.2-3x FLAG dendritic immunostaining upon loss of ankyrin-B (Figure 6A-C). Nav1.2-3x FLAG immunostaining at the AIS remained unchanged, consistent with ankyrin-G-dependent localization of Nav1.2-3x FLAG at the AIS (Figure 6A, B, D) (Jenkins et al., 2015). While rescue of Cre-mediated deletion of ankyrin-B with WT ankyrin-B-GFP resulted in the re-appearance of membrane-associated Nav1.2-3x FLAG immunostaining specifically at the dendrites, as evidenced by the “railroad track-like” FLAG staining, addition of palmitoylation-dead AAAAA-ankyrin-B-GFP did not rescue loss of Nav1.2-3x FLAG immunostaining at the dendrites in *Ank2* neocortical neurons (Figure 6A-C). These data suggest that ankyrin-B palmitoylation is required for proper dendritic localization of Nav1.2 in neocortical pyramidal neurons.

**Figure 6.**
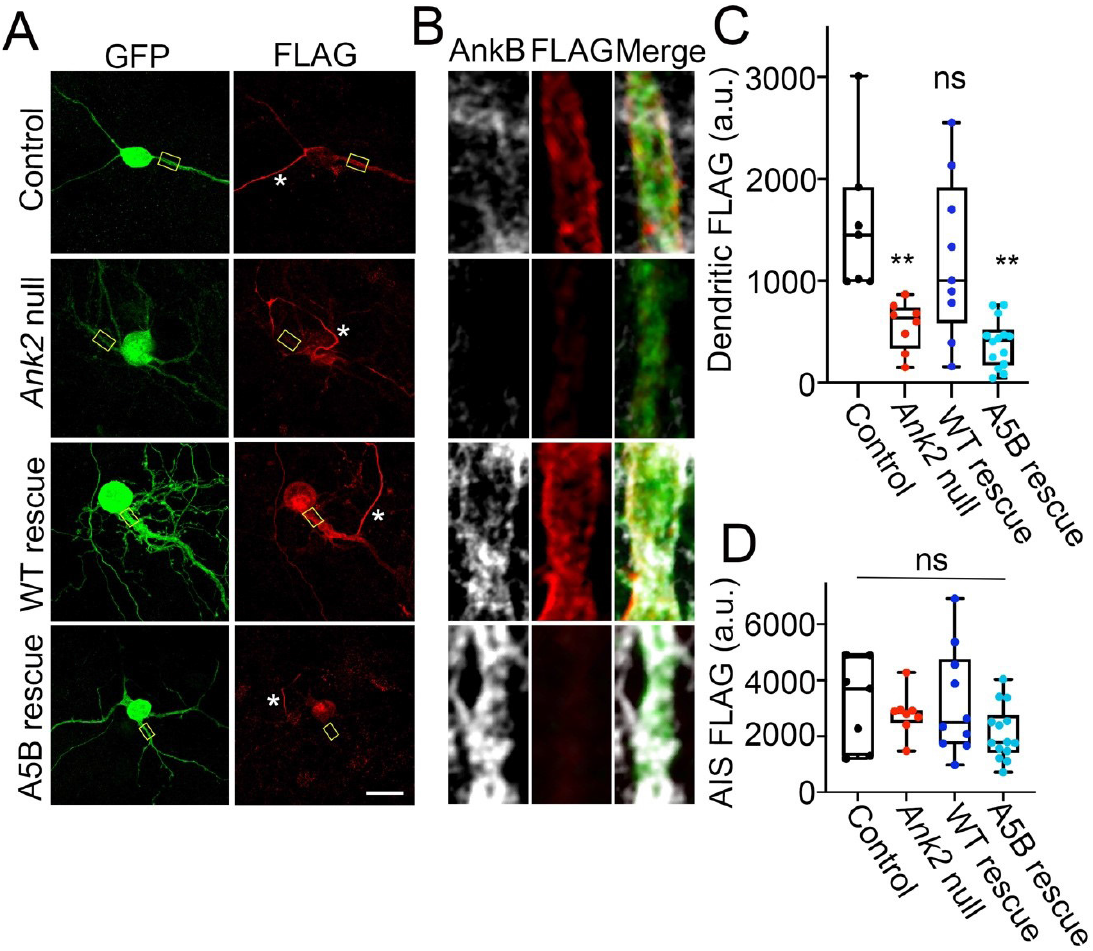
Loss of ankyrin-B palmitoylation prevents Nav1.2 from localizing properly at dendritic membranes of cultured neocortical neurons. **A:** Representative collapsed Z-stacks of DIV21 cultured floxed Ank2 cortical neurons transfected at DIV2 with Nav1.2-3x FLAG (Control) (Top), Cre-2A-BFP and Nav1.2-3x FLAG to generate total ankyrin-B-null neurons (Second from top), or Cre-2A-BFP, Nav1.2-3x FLAG, and indicated GFP rescue constructs (Bottom). Stars denote axon initial segment. Scale bars 20μm. Neurons stained with anti-GFP shown in green, anti-FLAG shown in red, and anti-ankyrin-B shown in far red. **B:** Zoomed-in single-stack images of selected dendritic compartment from A (denoted by yellow square). Anti-FLAG is shown in red, and anti-ankyrin-B is shown in far red. AAAAA-ankyrin-B-cannot rescue the loss of dendritic membrane localization of Nav1.2-3x FLAG induced by Cre-mediated deletion of ankyrin-B, compared to WT ankyrin-B-GFP. **C:** Quantification of mean fluorescence intensity of dendritic Nav1.2-3x FLAG from A,B. **P<0.01 relative to Control; one way ANOVA with Tukey’s post-hoc test, N=7-14 for each group across three independent replicates. **D:** Quantification of mean fluorescence intensity of AIS Nav1.2-3x FLAG from A,B. There is no change in AIS Nav1.2-3x FLAG localization for any of the conditions. NS (not significant) by one way ANOVA with Tukey’s post-hoc test, N=7-14 for each group across three independent replicates.

### Palmitoylation-dead AAAAA-ankyrin-B cannot properly localize to dendritic membranes of neocortical pyramidal neurons

We reasoned that the inability of palmitoylation-dead ankyrin-B to function in scaffolding Nav1.2 at dendritic membranes may be because palmitoylation-dead ankyrin-B is unable to properly target to dendritic membranes itself. To investigate whether dendritic localization of ankyrin-B is altered when ankyrin-B is unable to be palmitoylated, we first validated proper dendritic membrane localization of ankyrin-B as observed previously (Figure 6A, B) (Nelson et al., 2022). As expected, we observed ankyrin-B membrane localization at the dendritic plasma membrane, as evidenced by the “railroad-track” like appearance of ankyrin-B immunostaining in BFP-transfected control neurons (Figure 7A, B). We confirmed the efficiency of our knockout by verifying that dendritic membrane ankyrin-B localization was significantly reduced in *Ank2^flox/ flox^* neurons transfected with Cre-2A-BFP (*Ank2* null) compared to BFP-transfected (wild-type) control neurons (Figure 7A, B). Rescue of Cre-mediated loss of ankyrin-B with WT 220-kDa ankyrin-B-GFP resulted in the re-appearance of membrane-associated ankyrin-B-GFP staining at the dendrite, as shown by the two green GFP peaks flanking the soluble BFP peak in the plot profile (Figure 7C, D). Strikingly, rescue of Cre-mediated loss of ankyrin-B with palmitoylation-dead AAAAA-ankyrin-B-GFP resulted in loss of membrane-association of ankyrin-B, as demonstrated by the lack of “railroad-track” like appearance of AAAAA-ankyrin-B-GFP and as shown by the overlapping green GFP and blue soluble BFP peaks (Figure 7C, D). These data suggest palmitoylation-dead ankyrin-B is not able to associate with dendritic membranes and is instead mainly distributed in the dendrite cytoplasm. This phenotype is consistent with the fate of palmitoylation-dead neuronal ankyrin-G, C70A ankyrin-G, which is unable to associate specifically with the AIS membrane and instead distributes in a non-polarized fashion throughout the cytoplasm, rendering it nonfunctional and thus incapable of clustering binding partners such as neurofascin (Tseng et al., 2015). These data highlight that palmitoylation is required for ankyrin-B dendritic targeting, which is critical for ankyrin-B function in scaffolding Nav1.2 in neocortical pyramidal neurons.

**Figure 7.**
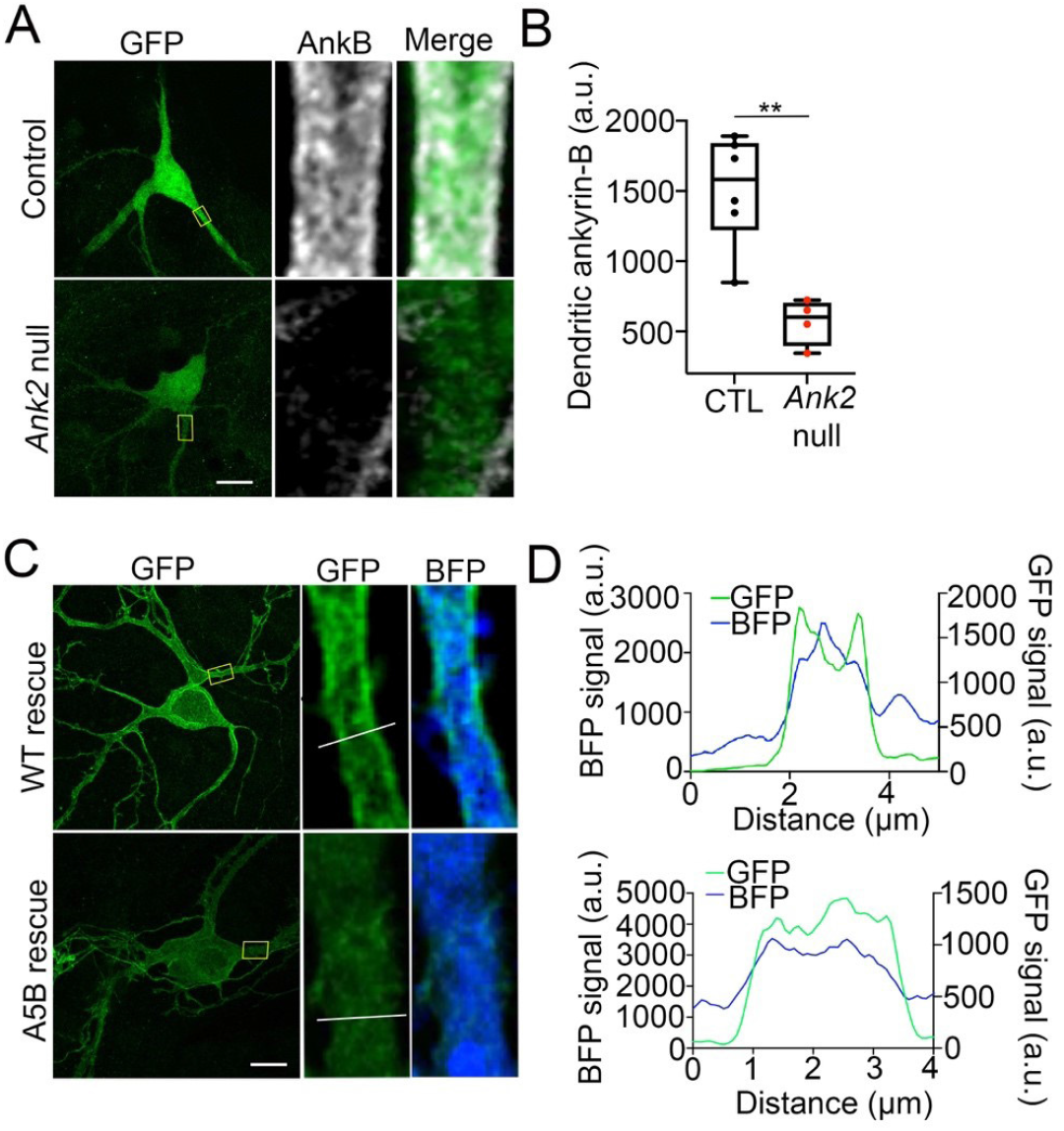
Palmitoylation-dead ankyrin-B is unable to target to dendritic membranes in neocortical pyramidal neurons. **A:** Representative collapsed Z-stack (left) and zoomed-in single stack images (right) of DIV21 cultured floxed Ank2 cortical neurons transfected at DIV2 with soluble GFP and BFP (top) or soluble GFP and Cre-2A-BFP (bottom) to knock-out ankyrin-B. Boxes denote zoomed-in section of the dendrite. Scale bars 20 μm. Neurons stained with anti-GFP shown in green and anti-ankyrin-B shown in far red. **B:** Quantification of mean fluorescence intensity of dendritic ankyrin-B from A. P**<0.01 relative to Control; student’s t-test (unpaired), N=4-6 neurons for each group across three independent replicates. **C:** Representative collapsed Z-stack (left) and zoomed-in single stack images (right) of DIV21 cultured floxed Ank2 cortical neurons transfected at DIV2 both transfected with Cre-2A-BFP to knock-out ankyrin-B and either rescued with WT ankyrin-B-GFP (top) palmitoylation-dead AAAAA-ankyrin-B-GFP (bottom) to assess dendritic membrane localization of ankyrin-B. Yellow boxes denote zoomed-in section of the dendrite. White line denotes region of interest across the dendrite used to generate plot profiles outlining membrane versus cytoplasmic ankyrin-B-GFP staining. Scale bars 20 μm. Neurons stained with anti-GFP shown in green. Blue fluorescence comes from 2A peptide from Cre-2A-BFP, which fills the cell as soluble BFP. **D:** Plot profile outlining membrane versus cytoplasmic GFP (ankyrin-B) (green) and BFP (blue) staining across dendrite (region of interest denoted by white line on zoomed-in single stack). What appears as plasma-membrane localization of ankyrin-B at dendrites is demonstrated by the two green peaks flanking the soluble BFP peak. The two green peaks in plot profile representing membrane-localized ankyrin-B are present in WT ankyrin-B-GFP rescue but not in the palmitoylation-dead ankyrin-B-GFP rescue. N=3 independent replicates.

## DISCUSSION

Ankyrin-B’s functions as a membrane organizer at the axon have been well-characterized. At the axon, ankyrin-B links the dynein/dynactin motor complex to organelles to promote fast axonal cargo transport, scaffolds the cell adhesion molecule L1CAM at the distal axon to repress axonal branching, and assembles a distal axonal cytoskeleton with βII-spectrin and α-II spectrin that restricts ankyrin-G positioning at the AIS (Galiano et al., 2012;Lorenzo et al., 2014;Yang et al., 2019). Recently, ankyrin-B, itself highly enriched in dendrites, was revealed to localize the voltage-gated sodium channel Nav1.2 to dendritic membranes to facilitate dendritic excitability (Nelson et al., 2022). Thus, ankyrin-B plays important roles in ensuring neuronal connectivity, polarization, and excitability. Dysfunction in *ANK2* is highly implicated in ASD, yet it remains unclear how *ANK2*, independently or by converging with the high-risk ASD gene *SCN2A* or other ASD-associated genes, contributes to ASD etiology.

Given the strong association between *SCN2A* and *ANK2* in ASD, and the importance of ankyrin-B-mediated scaffolding of Nav1.2 at dendrites for proper dendritic and synaptic function, it is important to understand how ankyrin-B targets to the dendritic membrane to allow for Nav1.2 localization. Previous studies showed other members of the ankyrin family, like ankyrin-R and ankyrin-G, were S-palmitoylated and that ankyrin-G palmitoylation was required for proper ankyrin-G localization and function in polarized epithelial cells and in neurons (Staufenbiel, 1987;He et al., 2012;He et al., 2014;Tseng et al., 2015). Here, we extend those findings to show for the first time that ankyrin-B is also S-palmitoylated, and relies on palmitoylation for proper localization and function at dendritic membranes in neurons. We also show ankyrin-B palmitoylation does not affect ankyrin-B-mediated axonal cargo transport.

Importantly, this work highlights that ankyrin-B utilizes distinct palmitoylation mechanisms compared to ankyrin-G, which is palmitoylated by functionally redundant zDHHC5 and zDHHC8 at a single cysteine residue, Cys70 (He et al., 2012;He et al., 2014). We show that ankyrin-B is capable of being palmitoylated by zDHHC17 in heterologous cells and further validated those findings by demonstrating zDHHC17 is a critical mediator of endogenous ankyrin-B palmitoylation in neurons. In heterologous cells, we observed zDHHC17 recognizes the zDABM domain of ankyrin-B to regulate both ankyrin-B expression levels and ankyrin-B palmitoylation. A previous study demonstrated the ANK domain of zDHHC17 recognizes a specific zDABM signature ((VIAP)(VIT)XXQP (where X is any amino acid)) in its substrate proteins in order to palmitoylate them (Lemonidis et al., 2014;Lemonidis et al., 2015;Lemonidis et al., 2017). Notably, the interaction between the ANK domain of zDHHC17 and the zDABM domain of ankyrin-B observed in this study was previously predicted by a peptide screen that identified the presence of the zDABM protein motif in ankyrins, suggesting our results are consistent with the conserved mechanism zDHHC17 utilizes to recognize and bind its substrates (Lemonidis et al., 2017). In line with the aforementioned study, we observed that the ability of ankyrin-B to recognize zDHHC17 is required for ankyrin-B palmitoylation, as demonstrated by the loss of ankyrin-B palmitoylation when the substrate binding site of zDHHC17 is ablated with the N100A mutation. Strikingly, loss of zDHHC17-ankyrin-B recognition also drastically reduced ankyrin-B protein levels. Although the enzymatic activity of zDHHC17 was required to regulate ankyrin-B palmitoylation, it was not sufficient to affect ankyrin-B expression levels. These data imply the observed zDHHC17-mediated increase in ankyrin-B protein expression is independent of palmitoylation, even though both protein expression and palmitoylation rely on recognition of ankyrin-B by zDHHC17.

The C60A and AAAAA-ankyrin-B-GFP mutants generated in this study to identify ankyrin-B palmitoylation sites also validated our findings that ankyrin-B protein expression was independent from ankyrin-B palmitoylation. While ankyrin-B palmitoylation was reduced by co-expression of either C60A or AAAAA-ankyrin-B-GFP with zDHHC17, protein expression of these mutant ankyrin-B plasmids was unchanged compared to WT ankyrin-B-GFP (Figure 4B, C, F, G). We reasoned this was likely because the zDABM in C60A and AAAAA-ankyrin-B-GFP remained intact, such that zDHHC17 is still capable of recognizing mutant ankyrin-B and thereby maintaining its expression. Our cycloheximide chase assays demonstrated that zDHHC17 does not stabilize ankyrin-B in our heterologous cell system with the experimental timepoints used for this assay. Thus, other potential palmitoylation-independent functions of zDHHC17 will need to be investigated to understand how zDHHC17 modulates ankyrin-B expression in heterologous cells.

Multiple studies have discussed the possibility of palmitoylation-independent functions of zDHHC17. Only half of the total number of proteins known to interact with zDHHC17 through their zDABM domain are known zDHHC17 substrates, suggesting that a major pool of zDHHC17 interactors rely on zDHHC17 for palmitoylation-independent functions (Lemonidis et al., 2015). Furthermore, the zDHHC17 orthologue zDHHC13 also has an ANK repeat domain capable of recognizing zDABM domains in proteins but is not capable of palmitoylating them (Lemonidis et al., 2014). The zDABM domain of peripheral membrane protein SNAP25 is capable of interacting with the ANK repeat domains of both zDHHC13 and zDHHC17, but is only capable of being palmitoylated by zDHHC17 and not zDHHC13 in heterologous cells (Lemonidis et al., 2014). These data suggest that although zDHHC17-zDABM binding is usually associated with palmitoylation, it can also serve additional substrate recruitment functions independent of zDHHC17 catalytic activity. The biological implications of ANK repeat-zDABM binding outside of zDHHC catalytic activity are only beginning to be uncovered. zDHHC17, by way of its ANK repeat domain, has been hypothesized to act as a hub for protein interaction networks. zDHHC17 is capable of forming large protein complexes by interacting with proteins involved in neurotransmission, neuronal development, trafficking, signal transduction, and transcriptional regulation, many of which have not been identified as palmitoylated substrates or zDHHC17-specific substrates (Butland et al., 2014). However, the functional implications of these interactions are currently unknown. zDHHC17 interacts with c-Jun N terminus kinase (JNK) to activate JNK and promote neuronal cell death in response to pathophysiological triggers like ischemic stroke (Yang and Cynader, 2011). zDHHC17 has been shown to promote the TrkA-tubulin complex, which regulates signal transmission in axon growth (Shi et al., 2015) Thus, our work is consistent with the ability of zDHHC17 to modulate substrates independently of its catalytic zDHHC domain, though this work is the first evidence to our knowledge that zDHHC17-zDABM binding directly regulates protein expression of a substrate in heterologous cells. However, what regulation of ankyrin-B protein levels by zDHHC17 means biologically and in what physiological context this finding is relevant will need to be further investigated. Consistent with our results in heterologous cells, we observed that a 95% reduction in Zdhhc17 mRNA in hippocampal neurons resulted in drastic loss of endogenous ankyrin-B palmitoylation. However, ankyrin-B protein levels remained unchanged in the absence of Zdhhc17 mRNA, suggesting that neurons may have other means of regulating ankyrin-B protein levels independently of zDHHC17. In light of recent work demonstrating zDHHC17 is localized to the somatic Golgi (Niu et al., 2020), ankyrin-B, which is highly localized to dendrites of mature neurons (Nelson et al., 2022), may not depend on zDHHC17 for maintenance of its protein pool out in the dendrites.

Functional studies with the palmitoylation-dead AAAAA-ankyrin-B-GFP demonstrated that palmitoylation regulates distinct functions of ankyrin-B. Palmitoylation of ankyrin-B was required for ankyrin-B to scaffold Nav1.2 at dendritic membranes of neonatal cortical pyramidal cells, but was not required for ankyrin-B to mediate axonal cargo transport of synaptotagmin-1. These data suggest there may be two pools of 220-kDa ankyrin-B in neurons, one palmitoylation-dependent pool at the dendrites that promotes Nav1.2 targeting and one palmitoylation-independent pool at the distal axon that promotes cargo transport. Consistent with the hypothesis that some palmitoylated substrates are locally palmitoylated (Philippe and Jenkins, 2019), somatic Golgi zDHHC17 may be conveniently located to palmitoylate the pool of 220-kDa ankyrin-B destined for dendrites at the soma before ankyrin-B is forward trafficked into dendrites, where it subsequently scaffolds Nav1.2. By contrast, zDHHC17 is specifically excluded from the axon in dorsal root ganglion neurons (Niu et al., 2020), which may explain why palmitoylation did not contribute to the axonal cargo function of 220-kDa ankyrin-B at the axon: 220-kDa ankyrin-B may simply not be palmitoylated at the axon due to the absence of zDHHC17 there. Interestingly, the pool of 220-kDa ankyrin-B that mediates cargo transport is targeted to vesicles and palmitoylation-independent, while the pool of 220-kDa ankyrin-B that scaffolds Nav1.2 channels at dendrites is plasma-membrane-associated and palmitoylation-dependent. Palmitoylation of the homologous ankyrin-G also drives ankyrin-G targeting at plasma membrane domains of epithelial cells and neurons (He et al., 2012;He et al., 2014). It may be that ankyrin palmitoylation defines the precise localization of plasma-membrane-associated pools of ankyrins, such as AIS-localized ankyrin-G and dendritic ankyrin-B, but other mechanisms, posttranslational or otherwise, may drive the specific targeting of intracellular membrane pools of ankyrins such as vesicular ankyrin-B, at the axon. If this is the case, then 440-kDa ankyrin-B, which is targeted to the axonal plasma membrane of neurons to scaffold the cell adhesion molecule L1CAM and repress axonal branching (Yang et al., 2019), may rely on palmitoylation for its axonal membrane targeting as well. Our observation that 440-kDa ankyrin-B is palmitoylated both in the brain and in neurons in Figure 1 is consistent with this hypothesis.

Despite the high homology shared between the ankyrin repeat domains of ankyrin-G and ankyrin-B, which harbor their respective palmitoylated cysteine sites, this study revealed non-conserved palmitoylation mechanisms between ankyrin family members. While ankyrin-G is palmitoylated by zDHHC5 and zDHHC8 at one cysteine site Cys70 (He et al., 2012;He et al., 2014), our study demonstrated that ankyrin-B is palmitoylated by zDHHC17 at multiple cysteine sites. This raises important questions about the biological implications of such distinct palmitoylation mechanisms, and may provide insight the long-standing question about how two highly homologous proteins like ankyrin-G and ankyrin-B target to distinct membrane sites and function in a non-overlapping manner. Studies have shown that divergent domains within ankyrins can confer specificity for ankyrin-B or ankyrin-G localization and function in various cell types (Mohler et al., 2002;He et al., 2013), though this has not yet been investigated in neurons. The highly divergent C-terminal domain confers specificity for ankyrin-B function at membranes of the sarcoplasmic reticulum (SR) in neonatal cardiomyocytes (Mohler et al., 2002). Additionally, the highly divergent short linker peptide between the ankyrin repeat domain and the ZU52-UPA module inhibits ankyrin-B binding to plasma membrane domains and drives intracellular membrane localization of ankyrin-B in epithelial cells (He et al., 2013). Given that no consensus has been reached on which domain(s) determine specificity for ankyrin-G or ankyrin-B localization, this work has brought forward the importance of considering palmitoylation as a potential mechanism underlying the distinct localization and functions of ankyrin-B and ankyrin-G.

In light of the recent discoveries highlighting ankyrin-B’s role in scaffolding Nav1.2 at dendrites to promote dendritic function (Nelson et al., 2022), it will be of interest to investigate the role of ankyrin-B palmitoylation for dendritic excitability and synaptic plasticity, especially as efforts to understand the convergent roles of ankyrin-B and Nav1.2 in the etiology of ASD continue. Our findings regarding the dendritic pool of ankyrin-B being palmitoylation-dependent and the vesicular pool of ankyrin-B at the distal axon being palmitoylation-independent highlights an opportunity to leverage palmitoylation as a drug target to specifically modulate the dendritic pool of ankyrin-B without affecting the vesicular pool of ankyrin-B at the axon, should the dendritic pool of ankyrin-B be implicated in ASD pathophysiology. It will also be important to investigate the role of Nav1.2 palmitoylation to understand how this may affect the formation and maintenance of the ankyrin-B/Nav1.2 complex at dendritic membranes.

## MATERIALS AND METHODS

### Antibodies

The antibodies used for western blotting in this study were chicken anti-GFP antibody (Abcam, 1:2000), rabbit anti-Flotillin-1 antibody (Cell Signaling Technologies, 1:1000), rabbit/mouse anti-HA antibody (Cell Signaling Technologies, 1:1000), and rabbit anti-ankyrin-B C-terminus (custom-made in-house, 1:1000). The specificity of the anti-ankyrin-B antibody has been shown previously by western blotting (Lorenzo et al., 2014). LiCor fluorescent secondary antibodies were used: IRDye 800CW donkey anti-rabbit and anti-chicken (for anti-ankyrin-B, anti-HA, anti-Flotillin-1) and IRDye 680RD donkey anti-mouse (for anti-HA) were diluted 1:10,000. The antibodies used for immunofluorescence studies were mouse anti-FLAG M2 (Millipore Sigma, 1:1000), chicken anti-GFP antibody (Abcam, 1:1000), and sheep anti-ankyrin-B C-terminus (custom-made, in-house, 1:1000). Alexa Fluor 488, 568, or 647 secondary antibodies (Fisher scientific) were used at 1:250.

### Expression Vectors

The panel (1-24) of mouse zDHHC cDNA constructs subcloned in a pEF-BOS HA backbone were kindly gifted to us by Dr. Masaki Fukata (National Institute of Physiological Sciences, Okazaki, Japan; (Fukata et al., 2004)). The zDHHA17 mutant generated by site-directed mutagenesis was kindly gifted to us by Dr. Luke Chamberlain (University of Strathclyde); (Locatelli et al., 2020)). The N100A zDHHC/A17 mutants were generated by our laboratory by site-directed mutagenesis using primers designed by Agilent QuickChange II XL primer design software and using QuickChange II XL site-directed mutagenesis kit (Agilent Technologies), and confirmed by DNA sequencing through Eurofins Genomics (Louisville, KY). Human ankyrin-B-GFP containing the small 220kD splice variant of ankyrin-B in a pEGFP-N1 vector was previously described (Lorenzo et al., 2014). All point mutations generated using the ankyrin-B-GFP cDNA construct were made using site-directed mutagenesis (QuickChange II XL, Agilent Technologies) and confirmed by DNA sequencing through Eurofins Genomics (Louisville, KY). Syt1-TdTomato was generously donated by Drs. Ronald Holz and Arun Anantharam (University of Michigan Medical School, Ann Arbor, MI). CAG-Cre-2A-BFP and CAG-BFP in a pLenti6-V5-DEST viral vector were previously used and described (Tseng et al., 2015).

### Animals

Wild type C57Bl/6J (Jackson Laboratories), floxed Zdhhc17 (on an FVB background (Sanders et al., 2016)), and floxed *Ank2* mice (previously described (Chang et al., 2014)) were housed in the Unit for Laboratory Animal Medicine at the University of Michigan. All procedures involving animals were performed in accordance with National Institutes of Health guidelines with approval by the Institutional Animal Care and Use Committee (IACUC) of the University of Michigan.

### Cell Culture and Transfections

HEK293T cells (gifted to us by Dr. Gareth Thomas, Temple University) were grown in Dulbecco’s modified Eagle’s high glucose medium with 10% fetal bovine serum, 1% penicillin, and 1% GlutaMax (ThermoFisher) at 37°C, 5% CO2. For Acyl-RAC experiments, HEK293T cells were plated in 10cm dishes at 80-90% confluency the day before transfection and transfected 16 hours later with 4 μg total DNA (2 μg of each plasmid) using Lipofectamine 3000 (Invitrogen) according to the manufacturer’s instructions.

### Neuronal Cultures, Transfections, and Viral transductions

Cortical neurons were dissociated from P0 floxed Zdhhc17 mouse pups and cultured were prepared as previous described (Nelson et al., 2018). At DIV2, neocortical cultures were transduced with β-galactose control adenovirus or Cre recombinase adenovirus, which are previously described (Davis et al., 2015;Brody et al., 2016;Brody et al., 2018). Cells were transduced in half of their original media for 6 hours before replacing with the remaining half of their original media and half of fresh, new media. Neurons were left to incubate for 10 days before being processed for RT-PCR or Acyl-RAC analysis.

Hippocampal neurons were dissociated from P0 *Ank2f/f* mouse pups and cultures were prepared as previously described (Nelson et al., 2018). For the axonal cargo transport imaging experiments, DIV3 hippocampal neurons were transfected with 1 μg of either CAG-BFP or CAG-Cre-2A-BFP cDNA in combination with 500 ng of the other plasmids (Synaptotagmin-1-TdTomato, WT ankyrin-B-GFP, AAAAA-ankyrin-B-GFP) using Lipofectamine 2000. Briefly, 1 μg or 1.5 μg of cDNA constructs were added to 200 μL of Neurobasal-A medium (Gibco) in an Eppendorf tube. In another tube, 3 μL or 4.5 μL, respectively, of Lipofectamine 2000 were added to 200 μL of Neurobasal-A medium. The two tubes were incubated separately in the hood for 5 minutes before merging, mixing well, and incubating for 20 more minutes. The neuronal growth media on the neurons was collected and saved at 37°C. The transfection mix was then added dropwise to the appropriate dishes of DIV3 neurons and incubated at 37°C for 1 h. The transfection mix was aspirated, cells were washed once with Neurobasal-A medium, and the original growth media was added back onto the neurons. Cells were maintained in culture for 48 h before live imaging.

For the dendritic ankyrin-B function and localization experiments, neocortical neurons were dissociated from P0 floxed *Ank2* mouse pups and cultures were prepared as previously described (Nelson et al., 2018). DIV2 neurons were transfected with 0.5 μg of each plasmid using Lipofectamine 2000 as described in the above paragraph.

### qRT-PCR

RNA was isolated from cultured hippocampal neurons in 6-well dishes (Thermo Fisher) using the RNeasy Mini Kit (QIAGEN). cDNA was prepared using the SuperScript IV First-Strand Synthesis Kit (Invitrogen). Quantitative RT-PCR on the mouse Zdhhc17 gene using primers spanning exons 1 and 2 (5’-ACCCGGAGGAAATCAAACCACAGA-3’ and 5’-TACATCGTAACCCGCTTCCACCAA-3’) and Sso/Advanced Universal SYBR green supermix (Fisher Scientific) was performed on CFX96 Real Time System (C1000 Touch Thermal Cycler, BioRad) under default conditions. Each sample was run in triplicates. Expression levels for mRNA were normalized to β-actin (5’-CATTGCTGACAGGATGCAGAAGG-3’ and 5’-TGCTGGAAGGTGGACAGTGAGG-3’).

### Acyl Resin Assisted Capture (Acyl-RAC)

Approximately 24h post-transfection, transfected HEK293T cells were lysed in “blocking buffer” containing 100 mM HEPES, 1 mM EDTA, 2.5% SDS, and 4% MMTS (Sigma), adjusted to pH 7.5 and sonicated. Samples were left to simultaneously heat and shake overnight at 40°C and 850 rpm in a Thermal Mixer II (Fisher Scientific). Samples were acetone precipitated with cold acetone (incubated in ice) to remove residual MMTS (previously described (Bouza et al., 2020)) and pellet was re-dissolved in 500 μL of “binding buffer” containing 100 mM HEPES, 1 mM EDTA, and 1% SDS, pH adjusted to 7.5, in Thermal Mixer II at 40°C and 850 rpm overnight. Protein samples were spun down in standard benchtop centrifuge at maximal speed for 5 minutes to pellet out any non-dissolved protein. Supernatant was then split into three 1.5 mL Eppendorf tubes, one containing 40 μL of unmanipulated starting material, one containing 200 μL of sample for hydroxylamine treatment (“+HA”), and one containing 200 μL of sample for NaCl treatment (“-HA”). 40 μL (1:1) of 5x SDS-PAGE buffer (5% wt/vol SDS, 25% wt/vol sucrose, 50 mM Tris pH 8, 5 mM EDTA, and bromophenol blue) supplemented with 100 mM DTT (Gold Biotechology) was added to the unmanipulated starting material and placed at 65°C for 10 minutes. 50 μL of 1:1 slurry of pre-activated thiopropyl-Sepharose 6b beads (GE, discontinued) were added to the “+HA” and “-HA” samples (previously described (Bouza et al., 2020)). 40 μL of freshly prepared 2 M hydroxylamine (HA) (Sigma), adjusted to pH 7.5, were added only to the “+HA” designated sample. 40 μL of 2 M NaCl were added to the “-HA” designated sample. Samples treated with HA and beads were left to incubate while rotating at room temperature for 2.5 h before being spun down at 5000 x g for 1 minute and washed 3x with “binding buffer”, each time discarding the supernatant and recovering the beads. 40 μL of 5x SDS-PAGE/DTT buffer were used to elute palmitoylated proteins off of the beads and heated for 10 minutes at 65°C. Samples were analyzed by western blotting.

For assessing palmitoylation of proteins from brain tissue or from cultured hippocampal neurons, after overnight MMTS block, samples were transferred to a Slide-A-Lyzer Dialysis Cassette (10,000 MWCO) (Thermo Scientific) for overnight buffer exchange with “Binding buffer”. Buffer exchanged samples were recovered from the dialysis cassette and the assay continued as described in the previous paragraph.

### Mass Spectrometry (LC-MS/MS)

After subjecting lysates to the Acyl-RAC assay and ensuring sufficient pulldown by western blotting (Supplementary Fig. 3A), the sample was submitted to the University of Michigan Mass Spectrometry-Based Proteomics Resource Facility (Department of Pathology). There, the thiopropyl sepharose beads (washed without SDS for optimal mass spectrometry analysis) were re-suspended in 50 μL of 8M urea/0.1M ammonium bicarbonate buffer (pH~8). Cysteines (including those holding the previously palmitoylated proteins to the beads) were reduced by adding 50 μL of 10 mM DTT and incubating at 45°C for 30 min. Samples were cooled to room temperature and cysteines were alkylated using 65 mM 2-Chloroacetamide in the dark for 30 minutes at room temperature. Upon diluting the urea to a final concentration of <1 M, sample was incubated with 1 μg sequencing grade, modified trypsin at 37°C overnight. Digestion was stopped by acidification and peptides were desalted using SepPak C18 cartridges using manufacturer’s protocol (Waters). Samples were completely dried using vacufuge. To perform the mass spectrometry, trypsinized peptides were dissolved in 9 μL of 0.1% formic acid/2% acetonitrile solution. Two μls of the resulting peptide solution were resolved on a nano-capillary reverse phase column (Acclaim PepMap C18, 2 micron, 50 cm, ThermoScientific) using a 0.1% formic acid/ acetonitrile gradient at 300 nl/min over a period of 180 min. Eluent was directly introduced into Orbitrap Fusion Tribrid mass spectrometer (Thermo Scientific, San Jose CA) using an EasySpray source. MS1 scans were acquired at 60K resolution (AGC target=3×106; max IT=50 ms). Data-dependent collision-induced dissociation MS/MS spectra were acquired on 20 most abundant ions following each MS1 scan (NCE ~28%; AGC target 1×105; max IT 45 ms). Proteins were identified by searching the data against Uniprot protein database using Proteome Discoverer (v2.4, Thermo Scientific). Search parameters included MS1 mass tolerance of 10 ppm and fragment tolerance of 0.2 Da. Two missed cleavages were allowed; carbamidomethylation of cysteines (+ 57.021 Da), oxidation of methionine (+ 15.995 Da), deamidation of aspergine and glutamine (+ 0.984 Da), and palmitoylation (+ 238.23 Da) were all considered variable modifications. False discovery rate (FDR) was determined using Percolator and proteins/peptides with an FDR of ≤1% (high confidence) were retained for further analysis.

### Western Blotting

Protein lysates, following Acyl-RAC or otherwise, were separated by 3.5-17% gradient gel in 1x Tris buffer, pH 7.4 (40 mM Tris, 20 mM NaOAc, and 2 mM NaEDTA) and 0.2% SDS at 175 Volts for ~4-5 hours. Transfer to a nitrocellulose membrane was performed overnight at 300 mAmps, 4°C in 0.5x Tris buffer and 0.01% SDS. Once transferred, membranes were blocked with 5% Bovine Serum Albumin (BSA) in 1x TBS and rotating overnight at 4°C with primary antibodies diluted at appropriate dilution factor in blocking buffer (5% BSA, 0.1% tween in TBS). Membranes were then washed 3x for 10 minutes each with 1x TBS-T (TBS with 0.1% tween) and incubated for 1 h with LiCor fluorescent antibodies in blocking buffer. Membranes were washed 3x for 10 minutes each in 1x TBS-T before imaging on LiCor Odyssey Clx Imager. Immunoreactive signals were quantified using ImageJ.

### Live imaging of hippocampal neurons and image analyses

Live microscopy of neuronal cultures was performed 48 hrs following neuronal transfections using a Zeiss LSM 880 with a 63X NA1.4 Oil/DIC Plan-Apochromat objective with excitation achieved using 405, 488, and 561 nm lasers in fast Airyscan mode. A humidified and temperature-controlled chamber was used to maintain the transfected hippocampal cultures at 37°C and 5% CO2 in warm Physiological Saline Solution (130 mM NaCl, 4 mM KCl, 1.5 mM CaCl2, 1 mM MgCl2, 5 mM Glucose, 10 mM HEPES, adjusted to pH 7.4, and filter sterilized). Time-lapse images captured transfected hippocampal neurons in the mid-axon so as to avoid the axon initial segment at a rate 200 ms intervals in fast Airyscan mode. Images were Airyscan-processed using the Zeiss image analysis software (Zen Blue). Kymographs were then generated from each time-lapse image using the KymoToolBox plugin in ImageJ (National Institutes of Health, (Schindelin et al., 2012)), as previously described (Zala et al., 2013). Briefly, the axon was outlined and traced, from which a kymograph calibrated in space (x-axis in micrometers) and time (y-axis in seconds) was generated. Individual particle movement/trajectory was traced along the kymograph, which the KymoToolBox plugin analyzed in terms of directionality (red for retrograde, green for anterograde, and blue for stationary), directional velocities (μm/seconds), and directional run length for each traced particle.

### Fluorescence Labeling of Cortical Neurons and image acquisition

Dissociated cortical neurons at DIV21 (transfected DIV2) were fixed for 15 min at room temperature with 4% formaldehyde, followed by permeabilization with 0.2% triton in PBS for 10 mins at room temperature, and further blocked with blocking buffer (5% BSA, 0.2% Tween 20 in PBS) for one hour. Primary antibodies were diluted in blocking buffer and incubated at 4°C overnight. The next day, cells were washed three times for 10 min with washing buffer (0.2% Tween 20 in PBS) before incubating the cells with secondary antibodies (1:250) diluted in blocking buffer for 1 h at room temperature. Cells were washed three times for 10 min with washing buffer and mounted with prolong gold. Multi-color imaging was performed as previously described using a Zeiss 880 confocal microscope (Jenkins et al., 2015). All images were processed in Adobe Photoshop CS6 and quantified using GraphPad Prism 9.

### Statistical Analysis

Statistical analyses for Acyl-RAC experiments were performed with n≥3 for each experiment ((N100A) zDHHC/A17: n=4, Zdhhc17/Cre neurons, n=4, C60A: n=7, 120kDa: n=3, for AAAAA: n=5). For the Zdhhc17f/f neuron Acyl-RAC, the C60A ankyrin-B Acyl-RAC, and the 120-kDa ankyrin-B Acyl-RAC experiments, a two-tailed student’s t-test (unpaired) was performed. Data are represented as mean ± SEM. For the Acyl-RACs that included multiple comparisons ((N100A) zDHHC/A17 and AAAAA-ankyrin-B), a one-way ANOVA with multiple comparisons and Tukey’s post-hoc test was performed. For the cargo transport experiment, four independent experiments were performed. Statistical significance for axonal velocity and run length of Syt1 particles, which included 4-8 axons per condition (n=88-154 particles per condition) was determined with one-way ANOVA with Tukey’s post-hoc test. To determine statistical significance for percentage of stationary, reversing, or motile Syt1-Tdtomato-positive or ankyrin-B-GFP-positive particles, a two-way ANOVA with multiple comparisons and Tukey’s post-hoc test was performed. Statistical significance for axonal velocity and run length of ankyrin-B particles, which included 3-7 axons per condition (n=38-76 particles per condition) was determined with a two-tailed student’s t-test (unpaired). All cargo transport data are represented as mean ± SEM. The Nav1.2-3x FLAG microcopy experiment had an n of 7-14 neurons/ condition and was performed in three independent replicates. Mean fluorescence intensity of dendritic FLAG signal and mean fluorescence intensity of AIS FLAG signal were analyzed using one-way ANOVA with multiple comparisons and Tukey’s post-hoc test. Data are represented as mean ± SEM.

## ABBREVIATIONS USED

HEK: Human Embryonic Kidney
AIS: Axon Initial Segment
ASD: Autism Spectrum Disorder
VGSC: Voltage-Gated Sodium Channel
PAT: Palmitoyl Acyl Transferase
zDABM: zDHHC ankyrin-repeat binding motif
GFP: Green Fluorescent Protein
BFP: Blue Fluorescent Protein
Syt1: Synaptotagmin-1
WT: Wild-Type
A5B: AAAAA-ankyrin-B-GFP
MMTS: S-methyl thiomethanesulfonate
RAC: Resin Assisted Capture
IB: Immunoblotting
RT: Room temperature
HA: Hydroxylamine
DIV: Days in Vitro
LCMS/MS: Liquid Chromatography tandem Mass Spectrometry
A: Alanine
C: Cysteine

## ACKNOWLEDGEMENTS

The authors thank Dr. William Fuller (University of Glasgow) for gifting us with functionalized thiopropyl beads, as well as Dr. Matthew Brody (University of Michigan) for providing feedback and guidance on this manuscript. The authors acknowledge support from Dr. Venkatesha Basrur with the University of Michigan Mass Spectrometry-Based Proteomics Resource Facility in the UM Department of Pathology.

## AUTHOR CONTRIBUTIONS

Conceptualization and methodology: JMP, PMJ; Investigation, formal analysis, and writing (original draft): JMP, Writing (review and editing): JMP and PMJ; Funding: JMP, PMJ; Supervision and resources: PMJ.

## SUPPLEMENTARY MATERIALS

Figures S1 to S3

