## Supplemental Figures for "Ankyrin-B is lipid-modified by S-palmitoylation to promote dendritic membrane scaffolding of voltage-gated sodium channel Nav1.2 in neurons"

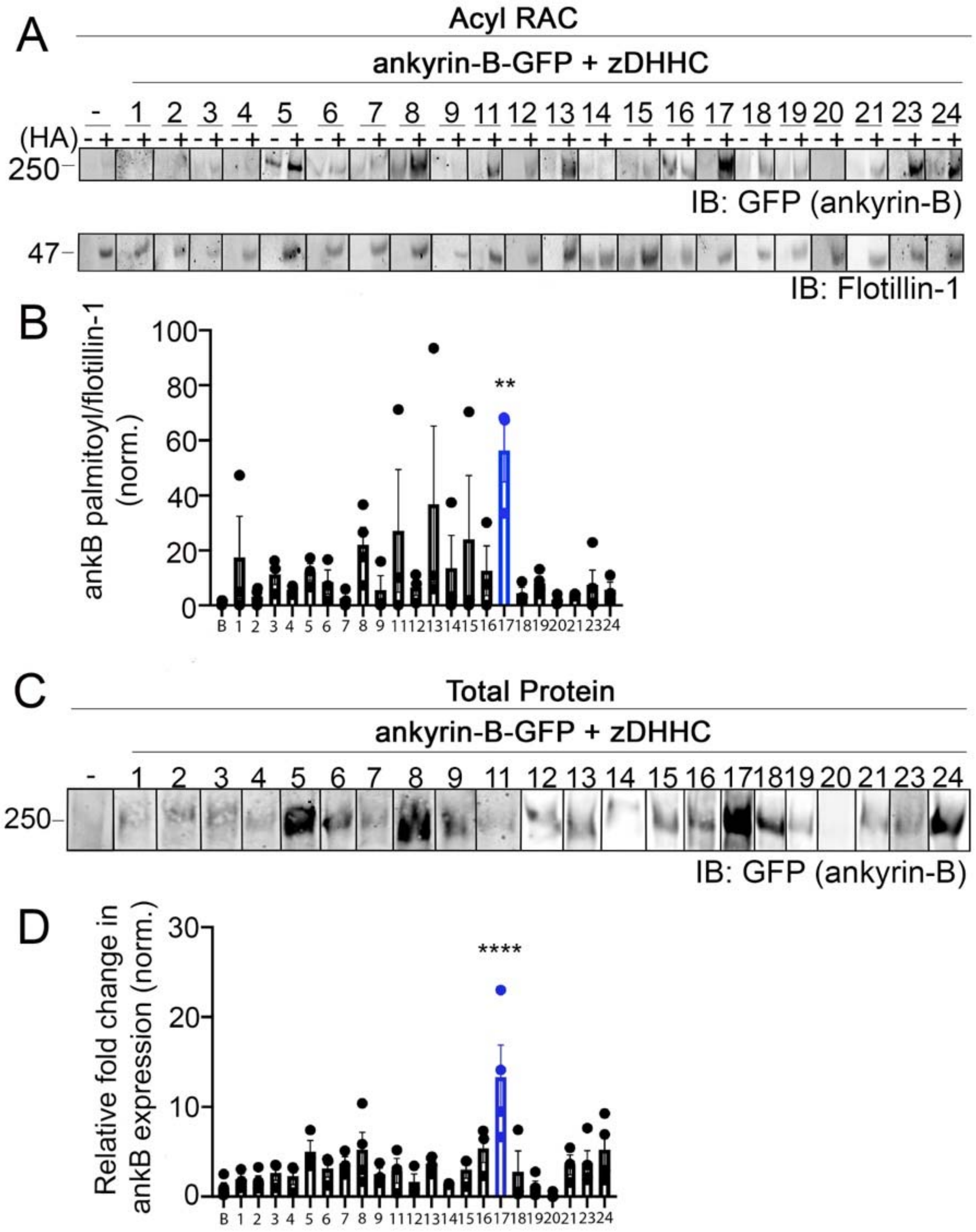


**Figure S1. zDHHC17 increases ankyrin-B palmitoylation and protein expression in HEK293T cells.**

**A.** Representative western blot showing ankyrin-B-GFP palmitoylation in the absence or presence of each individual zDHHC enzyme. Lysates from HEK cells transiently transfected with 220-kDa ankyrin-B-GFP alone (indicated as "-" in figure) or with each individual zDHHC enzyme were subjected to the Acyl-RAC assay to detect which zDHHC enzymes selectively enhance the palmitoylation of ankyrin-B-GFP compared to ankyrin-B alone. *S*-palmitoylation of ankyrin-B-GFP is detected using an antibody against GFP, as shown by the anti-GFP signal in '*+HA*' lane, compared to background signal in the negative control -*HA* lane. **B**. Quantification of *A*. zDHHC17 significantly enhances *S*-palmitoylation of ankyrin-B-GFP by approximately 91 fold compared to ankyrin-B-GFP alone (N=3). For each condition, the palmitoylation signal is calculated by subtracting the '*-HA*' lane signal from that of the '*+HA*' lane and normalizing to the flotillin signal from the '*+HA*' lane, and further normalizing to the average of ankyrin-B alone signal to get the relative fold change in ankyrin-B-GFP palmitoylation. Significance ($p = 0.0012$) was determined using an ordinary one-way ANOVA and Tukey's post-hoc multiple comparisons test. **C.** Representative western blot showing ankyrin-B-GFP total protein expression in the absence or presence of each individual zDHHC enzyme. **D**. Quantification of *C*. zDHHC17 significantly increases the protein expression of ankyrin-B-GFP by approximately 13 fold compared to ankyrin-B alone, while none of the other zDHHC enzymes significantly increase ankyrin-B-GFP expression (n=3), ****$p<0.0001$ (one-way ANOVA, Tukey's post-hoc).

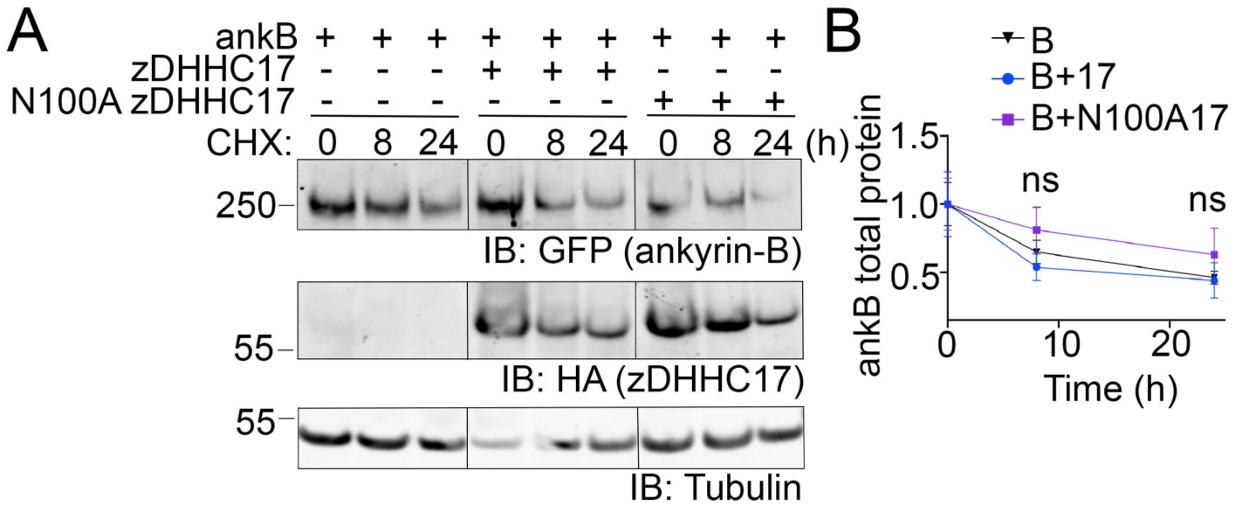


**Figure S2. zDHHC17 does not alter protein stability of ankyrin-B in heterologous cells by the cycloheximide (CHX) chase assay.**

**A.** Representative western blot showing ankyrin-B protein expression alone or in the presence of WT zDHHC17 or N100A zDHHC17 after 8 and 24 hours of cycloheximide treatment. HEK293T cells transiently transfected with 220-kDa ankyrin-B-GFP alone or with WT zDHHC17 or N100A zDHHC17 were treated with 100 µg/mL CHX and lysates were collected after 8 hours and 24 hours of CHX treatment before western blotting. Ankyrin-B expression is detected using an antibody against GFP. Absence or presence of zDHHC17 is detected using an antibody against HA. Tubulin is used as a loading control. For gel shown, all samples were run on the same gel to account for difference in baseline protein level; black line delineates spliced portion of the gel containing a condition not relevant to this figure. **B.** Quantification of *A*. Co-expression of zDHHC17 or N100A zDHHC17 does not alter the stability of ankyrin-B, compared to ankyrin-B in the absence of zDHHC17. For each condition, each time point is normalized to tubulin, and further normalized to Time 0h. Results are from N=6 for each condition; not significant (ns) by multiple unpaired *t-test*.

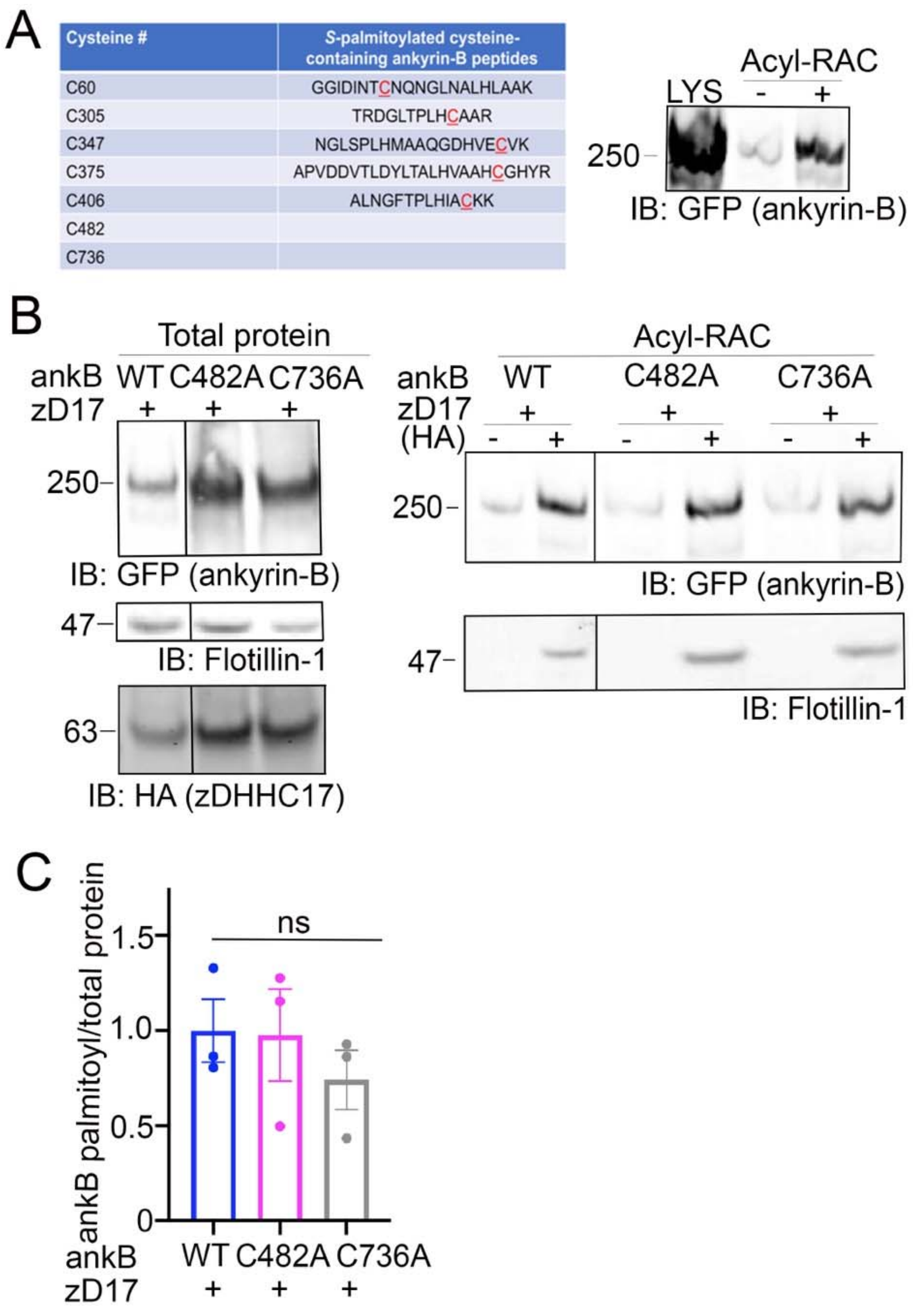


**Figure S3. Cys482 and Cys736 in ankyrin-B are not *S*-palmitoylated.**

**A.** Five cysteine-containing peptides were identified by mass spectrometry from HEK293T cells co-transfected with ankyrin-B-GFP and zDHHC17 processed for the Acyl-RAC assay. These cysteine-containing peptides corresponded to Cys60, Cys305, Cys347, Cys375, and Cys406 in ankyrin-B. Western blot demonstrates efficiency of the Acyl-RAC assay for sample containing overexpressed ankyrin-B-GFP and zDHHC17-HA submitted for mass spectrometry analysis. **B.** Validation that the two cysteines, Cys482 and Cys736 not identified as palmitoylated in the mass spectrometry analysis were indeed not palmitoylated by Acyl-RAC. Western blot shows total protein levels of ankyrin-B-GFP (*left*) and palmitoylation levels of ankyrin-B-GFP (*right*) from lysates of HEK293T cells transiently co-transfected with WT ankyrin-B-GFP, C482A ankyrin-B-GFP, or C736A ankyrin-B-GFP and zDHHC17 processed for the Acyl-RAC assay. No change in the palmitoylation levels of C482A or C736A ankyrin-B-GFP were observed compared to WT ankyrin-B-GFP, as evidenced by the unchanged level of GFP signal in the '+HA' lane for these two mutants compared to WT ankyrin-B-GFP. All conditions shown are run on the same blot; black line delineates spliced portion of the gel containing a condition not relevant to this figure. **C.** Quantified ankyrin-B-GFP *S*-palmitoylation levels normalized to total ankyrin-B protein levels for each condition, relative to palmitoylation levels of WT ankyrin-B-GFP co-expressed with zDHHC17. Data from N=3 independent replicates per condition from *B*. ns; one-way ANOVA, Tukey's post-hoc test.
